# The transcription factors TFEB and TFE3 link the FLCN-AMPK signaling axis to innate immune response and pathogen resistance

**DOI:** 10.1101/463430

**Authors:** Leeanna El-Houjeiri, Elite Possik, Tarika Vijayaraghavan, Mathieu Paquette, José A Martina, Jalal M. Kazan, Eric H. Ma, Russell Jones, Paola Blanchette, Rosa Puertollano, Arnim Pause

**Author notes:** Equal contribution (Leeanna El-Houjeiri, Elite Possik). **Corresponding Author:** Arnim Pause.

## Abstract

TFEB and TFE3 are transcriptional regulators of the innate immune response, but the mechanisms regulating their activation upon pathogen infection are poorly elucidated. Using *C. elegans* and mammalian models, we report that the master metabolic modulator 5’-AMP-activated protein kinase (AMPK) and its negative regulator Folliculin (FLCN) act upstream of TFEB/TFE3 in the innate immune response, independently of the mTORC1 signaling pathway. In nematodes, loss of FLCN or overexpression of AMPK conferred pathogen resistance *via* activation of TFEB/TFE3-dependent antimicrobial genes, while ablation of total AMPK activity abolished this phenotype. Similarly, in mammalian cells, loss of FLCN or pharmacological activation of AMPK induced TFEB/TFE3-dependent pro-inflammatory cytokine expression. Importantly, a rapid reduction in cellular ATP levels in murine macrophages was observed upon lipopolysaccharide (LPS) treatment accompanied by an acute AMPK activation and TFEB nuclear localization. These results uncover an ancient, highly conserved and pharmacologically actionable mechanism coupling energy status with innate immunity.

## Introduction

Innate immune responses constitute the first line of defense against pathogenic infections in simple metazoans, invertebrates, and mammals [1-4]. While much effort has been put into elucidating the functions of downstream mediators of immune response including antimicrobial peptides, C-type lectins, cytokines and chemokines, less is known regarding how host cells recognize foreign infections and trigger the activation of transcription factors that coordinate the anti-microbial response. Among the few well-characterized transcription factors, NF-κB, was shown to be an important factor in controlling host defense gene expression, mediated through toll like receptor (TLR) and nucleotide-binding leucine-rich repeat containing (NLR) ligand pathways [5]. However, another under-appreciated host-defense transcription factor was recently identified in *Caenorhabditis elegans* (*C. elegans)*, which lacks the NF-κB pathway [6]. Using this model, HLH-30, the *C. elegans* ortholog of TFEB and TFE3, was identified as an important evolutionarily conserved transcriptional regulator of the host response to infection [6-8]. TFEB and TFE3 are basic helix–loop–helix leucine zipper transcription factors that multi-task in regulating a similar set of genes involved in lipid metabolism, autophagy, lysosomal biogenesis and stress response genes [9-13]. Several studies have reported a similar mechanisms underlying TFEB/TFE3 activation in response to nutrient deprivation and metabolic stress. In nutrient-rich environments, the kinases ERK2 and mTORC1 phosphorylate TFEB/TFE3 on specific serine residues and retain them in the cytoplasm in an inactive state [13-17]. The mTORC1-dependent phosphorylation of TFEB (S211) and TFE3 (S321) promotes binding to 14-3-3. It has been suggested that this interaction masks the Nuclear Localization Signal (NLS), thus inhibiting TFEB and TFE3 nuclear translocation [14, 16]. Conversely, under starvation, this repressive phosphorylation is lifted, resulting in their translocation to the nucleus and activation of their downstream transcriptional targets that encode components of the lysosomal biogenesis and autophagy pathways [13, 14, 16, 17]. Despite these remarkable similarities between TFEB and TFE3, it is still unclear whether these transcription factors have cooperative, complementary, or partially redundant roles under different environmental conditions. Importantly, in murine macrophages, both TFEB and TFE3 were shown to be activated and translocated to the nucleus upon pathogen infection or stimulation with TLR ligands, where they collaborate in mediating the transcriptional upregulation of several cytokines and chemokines involved in antimicrobial immune response [6, 18, 19]. This functional conservation of the TFEB/TFE3 pathway is further supported by a recent study showing that bacterial membrane pore-forming toxin induces cellular autophagy in an HLH-30-dependent manner in *C. elegans* [20]. However, the mechanisms by which nematode and mammalian TFEB/TFE3 are activated during infection are still poorly understood. Recently, TFEB activation was found to involve phospholipase C and protein kinase D pathways both in *C. elegans* and mammals upon pathogen infection [21]. Subsequent studies showed that lipopolysaccharide (LPS)-stimulated TFEB/TFE3 activation in murine macrophages induced cytokine production and secretion independent of mTORC1, but the specific pathway by which their activation was mediated was not elucidated [18].

Folliculin (FLCN) is a binding partner and negative regulator of 5’-AMP-activated protein kinase (AMPK) [22, 23], which was identified as a tumor suppressor protein responsible for the Birt-Hogg-Dubé (BHD) neoplastic syndrome in humans [24]. Importantly, the interaction of FLCN with AMPK is mediated by two homologous FLCN-binding proteins FNIP1 and 2 [22,23]. Pathogenic mutations from BHD patients lead to a loss of FNIP/AMPK binding pointing to the functional significance of this interaction in tumor suppression [22]. AMPK is a heterotrimeric enzyme, which monitors the energy status and maintains energy homeostasis under metabolic stress by activating catabolic processes and inhibiting anabolic pathways [25-27]. We have previously shown that loss of FLCN or expression of a FLCN mutant unable to bind FNIP/AMPK led to chronic AMPK activation, resulting in increased ATP levels through an elevated glycolytic flux, oxidative phosphorylation and autophagy [28-30]. Importantly, we have shown that loss of FLCN mediates resistance to oxidative stress, heat, anoxia, obesity, and hyperosmotic stresses *via* AMPK activation in *C. elegans* and mammalian models [28-31].

While a role for FLCN in regulating immune responses has not been reported, the functional role for AMPK in innate immunity seems to be context and cell-type dependent [32, 33]. In this study, we demonstrate a novel evolutionarily conserved pathogen resistance mechanism mediated by FLCN and AMPK via TFEB/TFE3. Specifically, we show that loss of *flcn-1* in *C. elegans*, which leads to chronic AMPK activation, enhances the HLH-30 nuclear translocation and induces the expression of *hlh-30*-dependent antimicrobial genes upon infection, mediating resistance to bacterial pathogens. Using RNA-seq, we show that many *hlh-30*- dependent antimicrobial genes are regulated by AMPK upon *S. aureus* infection. AMPK loss reduces HLH-30 nuclear translocation and abrogates the increased resistance of *flcn-1(ok975)* mutant animals to pathogens. Furthermore, we show that constitutive activation of AMPK *C. elegans* nematodes leads to an HLH-30-dependent increase in pathogen resistance, similar to what we observe upon loss of *flcn-1*. Importantly, we show that this pathway of regulation is evolutionarily conserved and that FLCN and AMPK regulate TFEB/TFE3-driven cytokine and inflammatory genes in mouse embryonic fibroblasts and macrophages. Overall our data suggest an essential role of the FLCN/AMPK axis in the regulation of host-defense response via TFEB/TFE3, highlighting a possible mechanism likely to contribute to tumor formation in BHD patients. Our findings also shed new light on the potential use of AMPK activators in the stimulation of the innate immune response and defense against pathogens.

## Results

### Loss of *flcn-1* in *C. elegans* increases the expression of anti-microbial genes and confers resistance to bacterial pathogens

To understand the physiological role of FLCN-1, we compared gene expression profiles of wild-type and *flcn-1(ok975)* mutant animals. Among differentially expressed genes, 243 transcripts were up-regulated in *flcn-1(ok975)* mutant animals compared to wild-type animals at basal level (Table S1) and were classified based on their biological functions (Table 1 and Tables S2 and S3). Genes associated with stress response, innate immune response, defense mechanisms and response to stimulus processes, including heat shock proteins, C-type lectins, lysozymes and cytochrome P450 genes, were induced in *flcn-1(ok975)* unstressed mutant animals compared to wild-type animals (Table 1, S1-3, Figure 1A). Selected genes were validated using RT-qPCR (Figure 1B and Table S3). On the other hand, 704 genes were shown to be downregulated in *flcn-1(ok975)* mutant animals (Table S4) and are involved in various processes that control proliferation and growth (Table S5). These results indicate that a differential gene expression might be providing advantage to the *flcn-1* mutant worms prior to stress or pathogen attacks. This is in accordance with our previously reported results where loss of *flcn-1(ok975)* conferred resistance to oxidative stress, heat stress, anoxia and hyperosmotic stress in *C. elegans* [28-31, 34]. Since it was demonstrated that the osmo-sensitive gene expression mimics the transcriptional profiles of pathogen infection [35], we compared the overlap between genes upregulated in *flcn-1(ok975)* mutant animals and genes induced by infection of *C. elegans* nematodes with pathogens [36, 37]. Indeed, we found a significant overlap of the transcriptome especially upon *Staphylococcus aureus* (*S. aureus*) (Figure S1A and Table S6) and *Pseudomonas aeruginosa* (*P*. *aeruginosa*) infection (Figure S1B and Table S7).

**Table 1:**
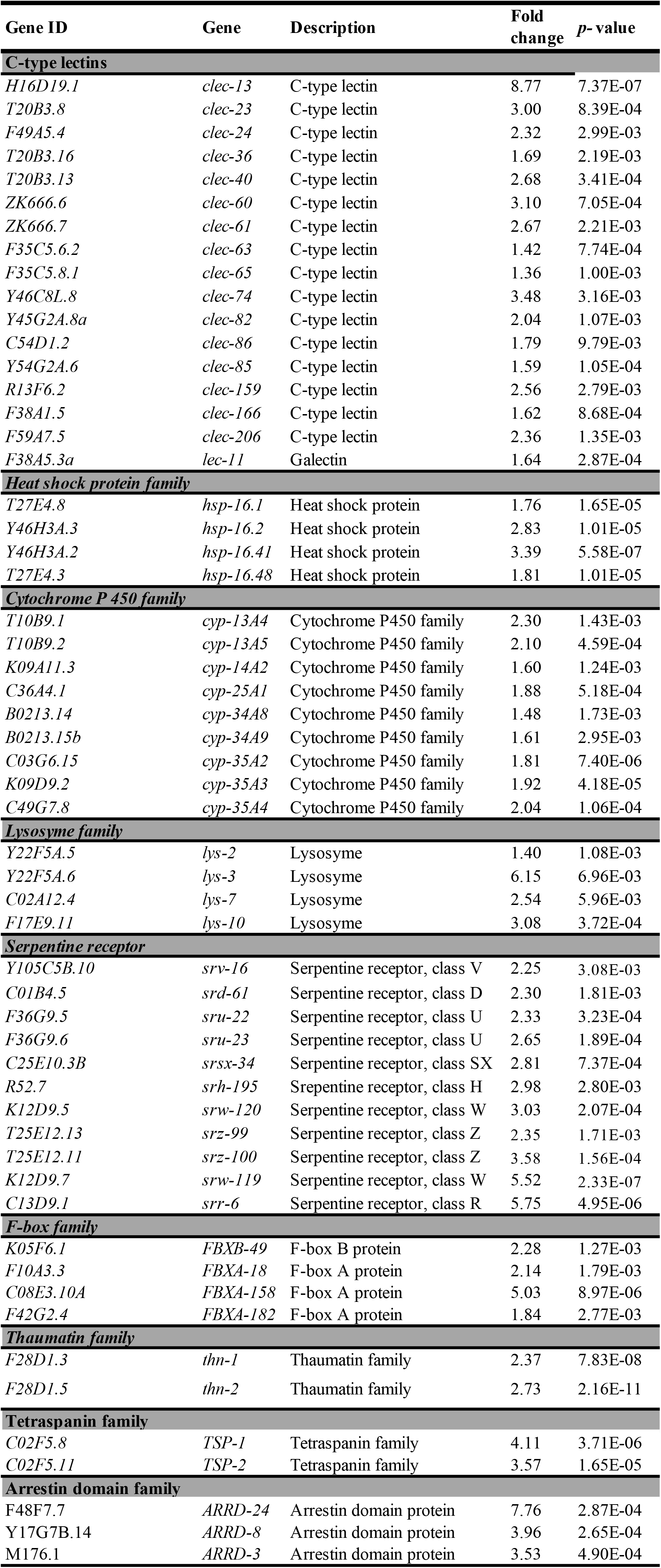
Genes classified according to family functions and upregulated in *flcn-1(ok975)* mutant animals.

**Fig 1:**
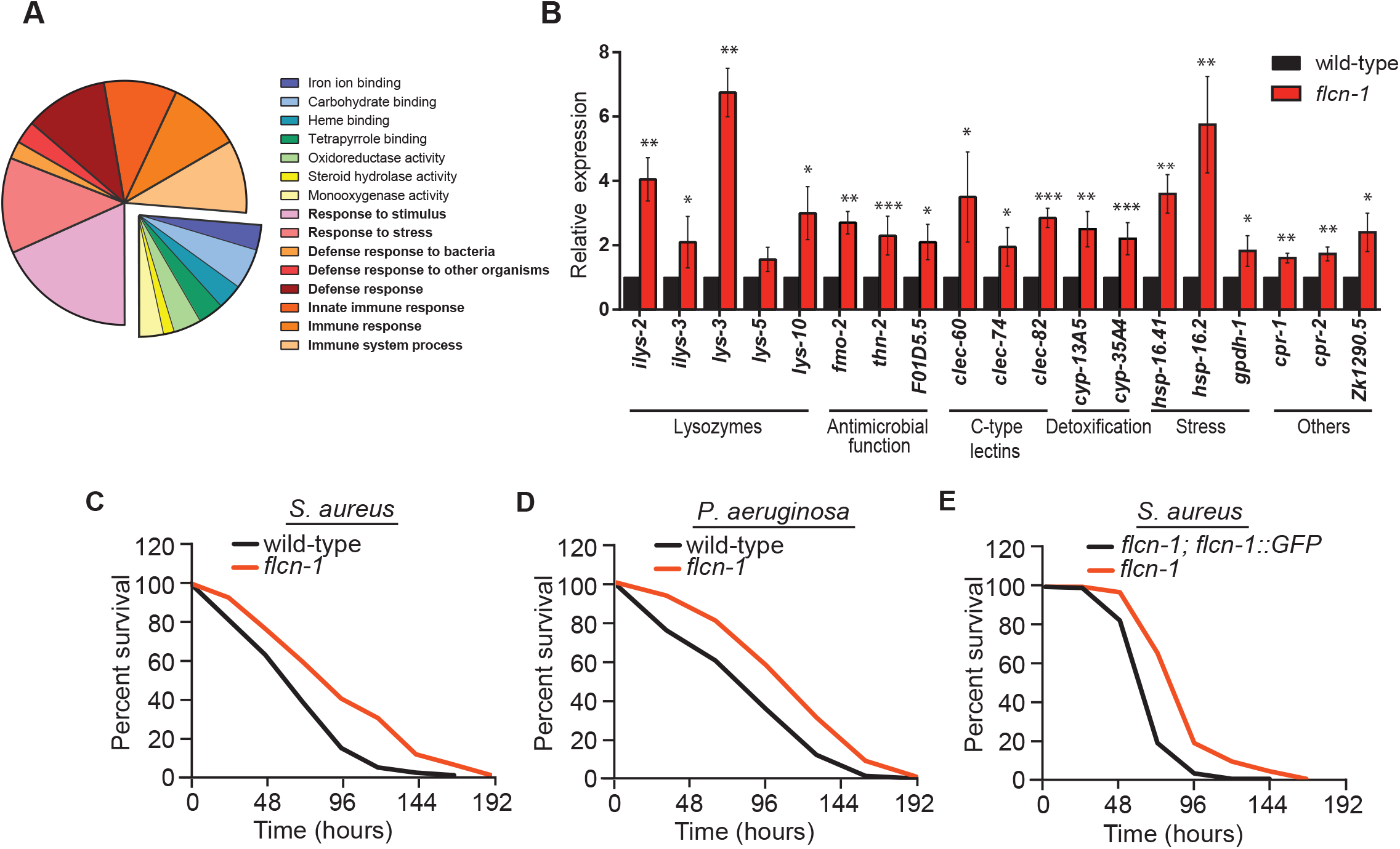
Loss of *flcn-1* increases the expression of antimicrobial genes and confers resistance to bacterial pathogens. (A) Pie chart of functional gene ontology analysis of the genes upregulated in *flcn-1(ok975)* mutant animals at basal level. (B) Relative mRNA expression of stress response and antimicrobial peptide genes in wild-type and *flcn-1* mutant animals. Data represents the average of three independent experiments, each done in triplicates ± SEM. Significance was determined using student’s t-test (^*^*p*<0.05, ^**^*p*<0.01, ^***^*p*<0.001). (C-E) Percent survival of indicated strains upon infection with *S. aureus* and *P. aeruginosa.* Refer to Tables S8A-B for details on number of animals utilized and number of repeats. The Statistical analysis was obtained using Mantel Cox test on the pooled curve.

Next, we asked whether *flcn-1(ok975)* mutant animals display enhanced resistance to pathogens. Strikingly, we found that the *flcn-1(ok975)* mutant animals are more resistant than wild-type animals to *S. aureus* and *P. aeruginosa* infection (Figure 1C-D, Tables S8A-B). These phenotypes were rescued using a transgenic *flcn-1* mutant animal re-expressing *flcn-1* (Figur 1E, Tables S8A-B). These results demonstrate an important role for *flcn-1* in the induction of antimicrobial peptides and stress response genes mediating the resistance to infection with bacterial pathogens.

### Loss of *flcn-1* increases pathogen resistance via HLH-30 activation

HLH-30, the worm ortholog of TFEB/TFE3, has been reported to modulate longevity and pathogen resistance in *C. elegans* through activation of autophagy and expression of antimicrobial genes [7, 12]. Importantly, we found a significant overlap between genes that were upregulated in *flcn-1(ok975)* mutant animals and downregulated in *hlh-30(tm1978)* mutant animals (Table S10) [6]. Thus, we asked whether HLH-30 is induced in *flcn-1* mutants using an *hlh-30*::GFP transgenic reporter strain [6, 7]. Upon infection with *S. aureus*, as shown in this study and others [6], HLH-30 translocated to the nucleus (Figure 2A). In particular, close to 40% of the wild-type animals displayed an HLH-30 nuclear localization after 20 min of infection with *S. aureus*. Importantly, we observed that the percentage of animals displaying a constitutive nuclear HLH-30 translocation in uninfected worms was significantly higher upon loss of *flcn-1* (Figure 2B, time 0). Strikingly, we show that upon *S. aureus* infection, the percentage of animals with HLH-30 nuclear translocation increased further in *flcn-1* mutant animals. Specifically, we found that after 20 min of infection with *S. aureus*, more than 80% of the *flcn-1* mutant animals displayed an HLH-30 nuclear localization in comparison to less than 40% for wild-type animals. Overall, this highlights an important role for HLH-30 in the increased pathogen resistance conferred by loss of *flcn-1* (Figure 2B).

**Fig 2:**
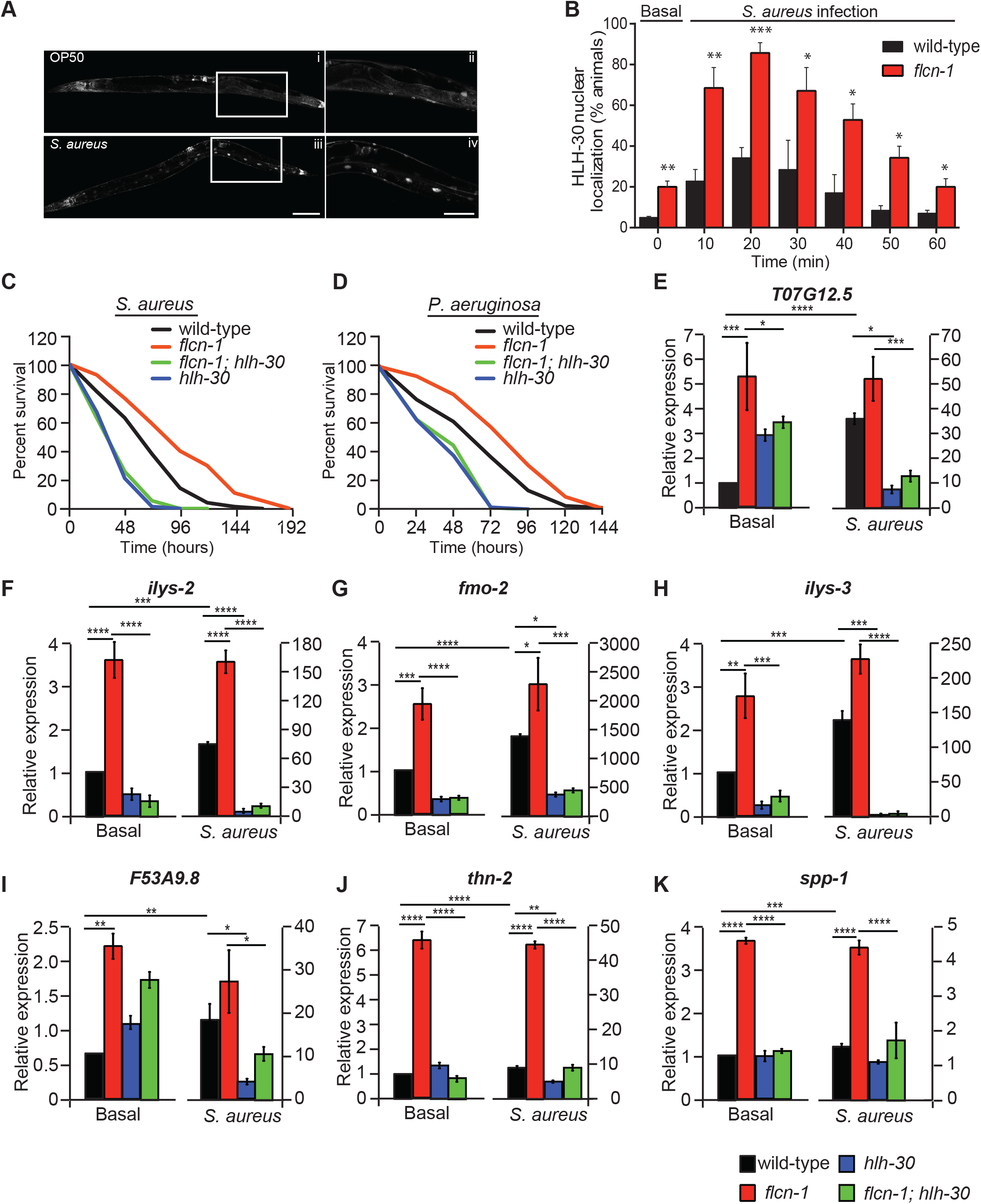
Loss of *flcn-1* increases pathogen resistance via HLH-30 activation. (A) Representative micrographs of HLH-30::GFP at basal level or after infection with *S. aureus* for 30 min. The signal is found in the nuclei of enterocytes, a cell-type in which lipids are stored in nematodes. Scale bars in i, iii and in ii, iv represent 100 μm and 50 μm respectively. (B) Percent animals showing HLH-30 nuclear translocation in *hlh-30p::hlh-30*::GFP and *flcn-1; hlh-30p::hlh-30*::GFP worm strains upon *S. aureus* infection for indicated time points to determine HLH-30 nuclear localization upon *flcn-1* loss at basal level (time 0) and upon *S. aureus* infection. Data represents the mean ± SEM from three independent repeats, n ≥ 30 animals/condition for every repeat. Significance was determined using student’s t-test (^*^*p*<0.05, ^**^*p*<0.01, ^***^*p*<0.001). (C, D) Percent survival of indicated worm strains upon infection with *S. aureus* and *P. aeruginosa*. Refer to Tables S8A-B for details on number of animals utilized and number of repeats. Statistics obtained using Mantel-Cox analysis on the pooled curve. (E-K) Relative mRNA expression of indicated target genes in wild-type, *flcn-1(ok975)*, *flcn-1(ok975); hlh-30 (tm1978)*, and *hlh-30 (tm1978)* animals at basal level and after treatment with *S. aureus* for 4 h. Data represents the average of three independent experiments done in triplicates ± SEM. Significance was determined using one-way ANOVA with the application of Bonferroni correction (^*^*p*<0.05, ^**^*p*<0.01, ^***^*p*<0.001).

To determine whether *hlh-30* is required for the increased survival of *flcn-1* mutant animals to pathogens, we generated a *flcn-1(ok975); hlh-30(tm1978)* double mutant strain. Importantly, loss of *hlh-30* significantly impaired the survival advantage upon both *S. aureus* (Figure 2C) and *P. aeruginosa* infections (Figure 2D) that was conferred by loss of *flcn-1*, demonstrating its involvement in pathogen resistance (Tables S8A-B). Accordingly, loss of *hlh-30* also suppressed the increased resistance of *flcn-1* to hyperosmotic stress [28] supporting that the adaptation to the two stresses requires a similar transcriptional profile dictated by HLH-30 (Figure S2).

To further assess the involvement of HLH-30 in the transcriptional response downstream of FLCN-1, we measured the gene expression of known HLH-30 target genes [6]. Using RT-qPCR, we found a significant upregulation in many *hlh-30*-dependent antimicrobial and infection-associated genes in uninfected *flcn-1* mutant worms (Figures 2E-K). Furthermore, after 4 h of infection with *S. aureus*, we show that loss of *hlh-30* strongly reduced the expression of antimicrobial peptide genes and infection-related genes in both wild-type and *flcn-1* mutant animals (Figures 2E-K), supporting a role for HLH-30 in the pathogen transcriptional signature downstream of *flcn-1*. Collectively, we found that loss of *flcn-1* activates the transcription of HLH-30 antimicrobial peptide genes at basal level, which is further induced upon *S. aureus* infection.

### The regulation of TFEB/TFE3 by FLCN is evolutionarily conserved through an mTOR independent mechanism

Because the role of HLH-30 in host defense is evolutionarily conserved [6], we tested whether the FLCN-HLH-30 axis that we uncovered in *C. elegans* is conserved from worms to mammals. Indeed, we observed that *Flcn* deletion in mouse embryonic fibroblasts (MEFs) promoted TFEB and TFE3 nuclear localization at basal levels compared to wild-type MEFs as detected by subcellular fractionation and immunofluorescence assays (Figure 3A-C). The difference in the cytosolic TFEB molecular weight can be attributed to the phosphorylation forms of TFEB [17]. Consequently, known TFEB and TFE3 targets were upregulated at the mRNA level upon *Flcn* deletion (Figure 3B), including genes involved in innate host response, such as IL-6. Addition of Torin1, a specific inhibitor of mTORC1 and mTORC2, induced TFEB nuclear localization in *Flcn* knockout (KO) MEFs to a higher extent than wild-type MEFs (Figure 3C), evoking an mTOR independent pathway. Moreover, loss of *Flcn* did not affect mTOR signaling, as measured by immunoblotting for the phosphorylated form of the S6 ribosomal protein (S6), a well-described mTORC1 downstream target (Figure 3D). In line with this, in *C. elegans*, inhibition of *let-363*, the *C. elegans* TOR homolog, increased the HLH-30 nuclear translocation at basal level similar to what has been previously reported [7] (Figure 3E). Importantly, loss of *flcn-1* further increased the HLH-30 nuclear translocation upon inhibition of *let-363* at basal level supporting a TOR-independent pathway governing HLH-30 regulation (Figure 3E). Moreover, infection with *S. aureus* increased to a similar extent the HLH-30 nuclear localization both in wild-type and *flcn-1 (ok975)* animals fed with *let-363* RNAi, presumably because the infection happens rapidly masking the effects of *let-363* RNAi on HLH-30 translocation (Figure 3E). These findings suggest that loss of FLCN drives HLH-30/TFEB/TFE3 nuclear localization through a mechanism distinct from the canonical mTOR pathway both in nematodes and mammalian cells.

**Fig 3:**
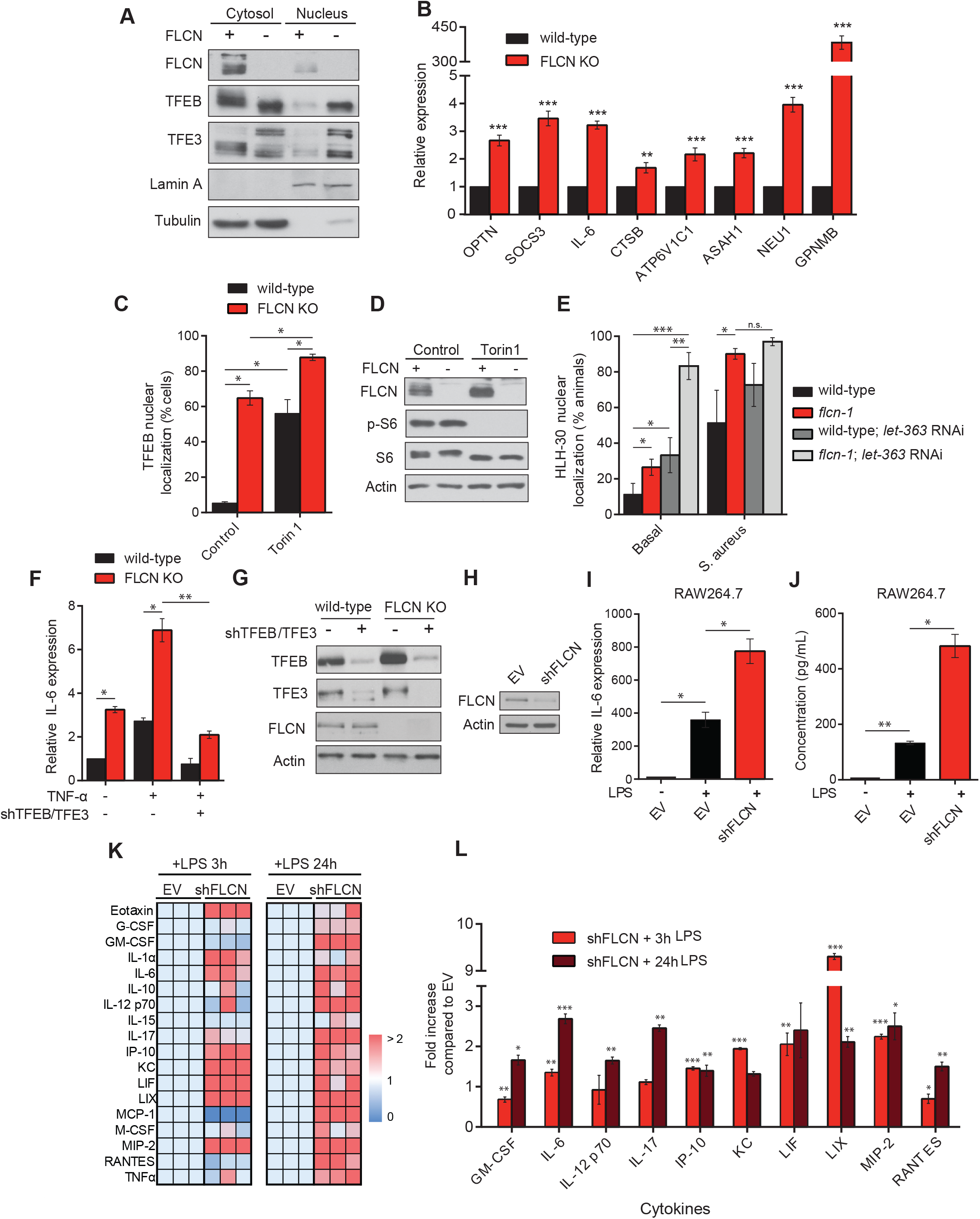
The regulation of TFEB/TFE3 by FLCN is evolutionarily conserved through mTOR independent mechanisms. (A) Immunoblot of isolated cytosolic-soluble fractions and nuclear fractions of wild-type and FLCN KO mouse embryonic fibroblasts (MEFs). (B) Relative mRNA levels measured by qRT-PCR of indicated genes in wild-type and FLCN KO MEFs. Data represent the average of three independent experiments done in triplicates ± SEM. Significance was determined using student’s t-test (^**^*p*<0.01, ^***^*p*<0.001). (C) Quantification of the percentage of cells showing TFEB nuclear staining treated with mTORC1 inhibitor; Torin1 (1 μM) for 2 h. Data represents the average of three independent experiments, each done in triplicates ± SEM. Significance was determined using one-way ANOVA with the application of Bonferroni correction (^*^*p*<0.05). (D) Immunoblot analysis of whole cell extracts with or without Torin1 (1 μM) for 2 h. (E) Percent animals showing HLH-30 nuclear translocation in indicated *hlh-30p::hlh-30::GFP* worm strains treated with or without *let-363* RNAi at basal level or upon *S. aureus* infection. Data represent the mean ± SEM with 3 independent repeats, n ≥ 30 animals/condition for every repeat. Significance was determined using one-way ANOVA with the application of the Bonferroni correction (^**^*p*<0.01, ^***^*p*<0.001). (F) Relative IL-6 mRNA levels measured by qRT-PCR in empty vector (EV) or shTFEB/TFE3-treated wild-type or FLCN KO MEFs, stimulated with or without 10 ng/ml TNF-α for 2 h. Data represents the average of three independent experiments, each done in triplicates ± SEM. Significance was determined using one-way ANOVA with the application of Bonferroni correction (^*^*p*<0.05, ^**^*p*<0.01). (G) Immunoblot analysis of wild-type and FLCN KO MEFs transfected with EV or shTFEB/TFE3. (H) Immunoblot analysis of RAW 264.7 cells transfected with EV or shFLCN. (I) Relative IL-6 mRNA levels measured by qRT-PCR in EV or shFLCN-treated RAW264.7 cells, stimulated with or without 1 μg/ml LPS for 3 h. (J) Quantification of IL-6 levels of conditions described in (I) using Mouse Protein Cytokine Array. Data represent the average of three independent experiments ± SEM. Significance was determined using one-way ANOVA with the application of Bonferroni correction (^*^*p*<0.05, ^**^*p*<0.01). (K) Hierarchical clustering of cytokine and chemokine secretion in the supernatant using Mouse Protein Cytokine Array in EV or shFLCN-treated RAW264.7 cells stimulated with 1 μg/ml LPS for 3 and 24 h. Each square in a column represents the average of triplicate experiments, and each column represents an independent replicate. Fold increase was normalized against EV and color-coded (dark red indicates 2 or more-fold increase, dark blue indicates no change). (L) Fold increase in cytokine and chemokine secretion levels as described in (J). Data represent the average of three independent experiments, each done in triplicates ± SEM. Significance was determined using student’s t-test in comparison to the EV stimulated with LPS for 3h and 24, respectively (^*^*p*<0.05, ^**^*p*<0.01, ^***^*p*<0.001).

To further assess whether the transcriptional up-regulation of cytokines and chemokines upon loss of *Flcn* was mediated by TFEB and TFE3, we knocked down their endogenous expression simultaneously using shTFEB and shTFE3 in wild-type and *Flcn* KO MEFs and determined the expression of IL-6 following TNFα stimulation (Figure 3F-G). Notably, we found that the significant induction of IL-6 mRNA levels upon TNF-α stimulation in both wild-type and *Flcn* KO MEFs was abrogated to levels observed in unstimulated cells upon knockdown of TFEB/TFE3 (Figure 3F). To confirm the observed effects in a relevant cellular system for innate immune response, we used RAW264.7 murine macrophages and reduced the endogenous expression of *Flcn* using shRNA-mediated knockdown approaches. Importantly, we show a significant increase in IL-6 production, at both mRNA (Figure 3H) and protein levels (Figure 3I), in FLCN KD macrophages compared to empty vector (EV) in response to LPS stimulation. To further assess the role of FLCN in inflammation and innate immune response, we determined the cytokine and chemokine secretion profiles in wild-type and shFLCN macrophages after 3h and 24h of LPS stimulation using mouse protein cytokine arrays. Notably, we show a significant and prominent increase in many cytokines in FLCN KD macrophages as compared to EV upon LPS stimulation (Figures 3J-K). These cytokines encompass key mediators of the inflammatory response.

### FLCN depletion in macrophages enhances their energy metabolism and phagocytic potential

Next, we investigated the metabolic consequences of FLCN depletion in RAW264.7 macrophages. We found that glucose consumption and lactate production were increased in FLCN KD macrophages compared to control macrophages (Figure 4A-B), and this was accompanied by an augmented extracellular acidification rate (ECAR) (Figure 4C-D) and oxygen consumption rate (OCR) (Figure 4E-F) at basal level and upon the sequential addition of oligomycin (an ATP synthase inhibitor), Carbonyl cyanide-p-trifluoromethoxyphenylhydrazone (FCCP) (for maximum respiratory capacity), followed by rotenone/antimycin A (to block mitochondrial electron transport). In line with these results, we also report an increase in ATP production in FLCN KD macrophages compared to controls (Figure 4G). Next, we investigated whether FLCN depletion enhances the phagocytic potential in macrophages (Figure 4I). Using pHrodo Red *S. aureus* Bioparticles, we report a 30% increase in phagocytic capacity of FLCN KD macrophages compared to control cells, as shown by the fold change in the mean florescence intensity (Figure 4I). To test whether this increase in phagocytic activity in FLCN KD macrophages is dependent on TFEB/TFE3 activation, we knocked down FLCN in TFEB/TFE3 DKO RAW macrophages (Figure 4H), and showed that the phagocytic activity of these cells decreased by almost 50% compared to FLCN depleted macrophages, upon stimulation with pHrodo Red *S. aureus* Bioparticles (Figure 4I). Taken together, we show that depletion of FLCN in macrophages prompts a metabolic transformation toward increased cellular bioenergetics, accompanied by an augmented TFEB/TFE3-dependent phagocytic capacity, which might further enhance the innate immune response.

**Fig 4:**
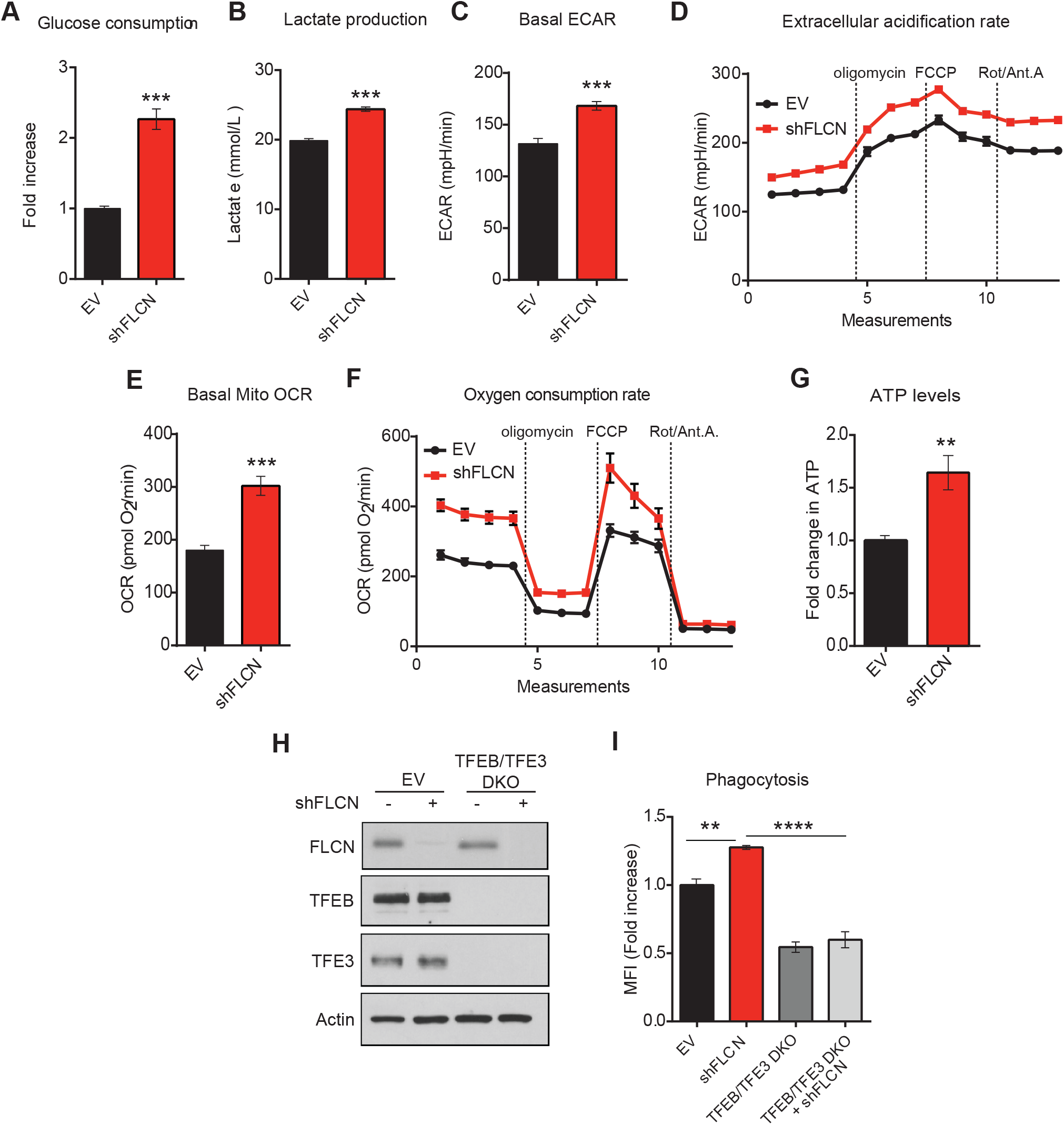
FLCN depletion in macrophages enhances their energy metabolism and phagocytic potential. (A) Glucose production and (B) lactate consumption levels measured using NOVA Bioanalysis flux analyzer in empty vector (EV) or shFLCN RAW264.7 at basal level. (C, D) Extracellular acidification rate (ECAR) and (E, F) oxygen consumption rate (OCR) of EV or shFLCN RAW264.7 at basal level as measured by Seahorse Bioscience XF96 extracellular flux analyzer. After establishing a baseline, oligomycin (10 μM), FCCP (15 μM), and rotenone/antimycin A (1 μM, and 10 μM, respectively) were added. (G) Fold change in ATP levels in EV or shFLCN RAW264.7 after 24 h of seeding as measured by CellTiter-Glo Luminescent Cell Viability Assay. Data represent the average of three independent experiments, each done in triplicates ± SEM. Significance was determined using student’s t-test (^**^*p*<0.01, ^***^*p*<0.001). (H) Immunoblot analysis of EV and TFEB/TFE3 DKO RAW264.7 cells transfected with EV or shFLCN. (I) Phagocytic activities of EV, TFEB/TFE3 DKO, and TFEB/TFE3 DKO shFLCN RAW264.7 cells measured using Red pHrodo *S.aureus* BioParticles by flow cytometry. Data represents the average of three independent experiments, each done in triplicates ± SEM. Significance was determined using one-way ANOVA with the application of Bonferroni correction (^**^*p*<0.01, ^****^*p*<0.0001).

### AMPK regulates HLH-30 activation and antimicrobial response upon infection with bacterial pathogens

Given that we have previously reported that loss of *flcn-1* leads to chronic AMPK activation, which increases resistance to energy [29] and hyperosmotic stresses [34] in nematodes, we tested whether *flcn-1* mutant animals confer pathogen resistance via AMPK-mediated regulation of HLH-30. Importantly, simultaneous loss of *aak-1* and *aak-2* (*C. elegans* orthologs of AMPK α1/α2) completely abolished the increased survival to both *S. aureus* and *P. aeruginosa* in wild-type and *flcn-1* mutant animals, demonstrating that this phenotype requires AMPK (Figures 5A-B and Tables S8A-B). Furthermore, transgenic overexpression of a constitutively active catalytic subunit of AMPK (*aak-2 oe*) in nematodes confers pathogen resistance similar to *flcn-1(ok975)* mutants, which is mostly dependent on HLH-30 (Figure 5C and Table S8 A). Moreover, loss of both AMPK catalytic subunits significantly reduced the nuclear translocation of HLH-30 upon *S. aureus* infection (Figure 5D). Additionally, we found that loss of *aak-2(ok524)* alone was insufficient to reduce the nuclear translocation of HLH-30 upon *S. aureus* infection, suggesting that complete abrogation of both AMPK catalytic activities is required for this phenotype (Figure S3B). To further elaborate the role of AMPK in pathogen response and specifically in the transcription of antimicrobial and stress response genes upon infection, we used RNA-seq technology to measure differential gene expression in wild-type and *aak-1(tm1944); aak-2(ok524)* mutant animals at basal level and after 4 h infection with *S. aureus* (Figure S3A, Tables S11-S15). We identified more than 800 genes induced upon *S. aureus* infection that are dependent on AMPK (Figure 5E and S3A, Tables S16-S17). Furthermore, we found a significant overlap of 112 genes down-regulated in *aak-1(tm1944); aak-2(ok524)* mutant animals and genes regulated by *hlh-30* upon *S. aureus* infection [6] (Figure 5F, Table S18). Gene ontology classification highlights important pathways regulated by AMPK during *S. aureus* infection, including defense response and stress response pathways (Figure 5G and Tables S16- S19). Using RT-qPCR, we validated several genes obtained by RNA-seq (Figures 5H-R), all of which have been reported to be involved in defense mechanisms against bacterial pathogens [3, 36]. Overall, these results indicate that AMPK regulates the nuclear translocation of HLH-30 and the HLH-30 driven antimicrobial response upon infection with bacterial pathogens.

**Fig 5:**
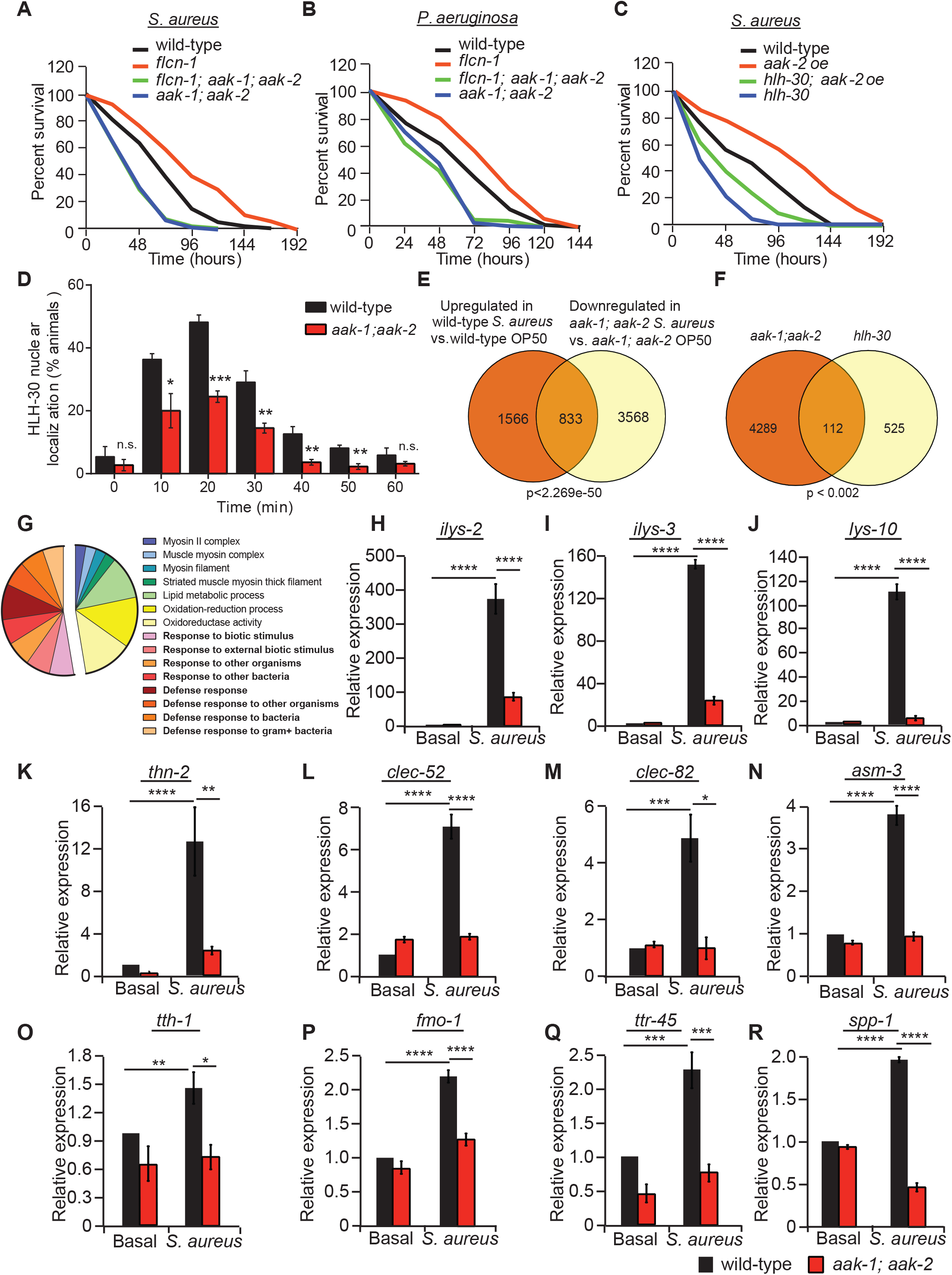
AMPK regulates HLH-30 activation and antimicrobial response upon infection with bacterial pathogens. (A-C) Percent survival of indicated worm strains upon infection with *S. aureus* or *P. aeruginosa*. Refer to Tables S8A-B for details on number of animals utilized and number of repeats. Statistics obtained by Mantel-Cox analysis on the pooled curve. (D) Percentage of animals showing HLH-30 nuclear translocation in indicated *hlh-30p::hlh-30::GFP* worm strains upon infection with *S. aureus* for the indicated amount of time. Data represent the mean ± SEM with 3 independent repeats, n ≥ 30 animals/condition for every repeat. Significance was determined using student’s t-test (^*^*p*<0.05, ^**^*p*<0.01, ^***^*p*<0.001). (E) Venn diagram of the overlapping set of genes between *S. aureus*-induced genes in wild-type animals and genes downregulated in *aak-1(tm1944); aak-2(ok524)* mutant animals upon infection. (F) Venn diagram and (G) pie chart of functional gene ontology analysis of AMPK-dependent genes obtained by the overlap analysis between genes downregulated in *aak-1(tm1944); aak-2(ok524)* mutant animals in comparison to wild-type animals upon *S. aureus* infection and the *hlh-30*-dependent list of genes published in (40). Comparisons were performed using the “compare two lists” online software and the significance was obtained using “nemates” software. (H-R) Relative mRNA levels measured by qRT-PCR of AMPK-dependent genes in wild-type and *aak-1(tm1944); aak-2(ok524)* mutant animals infected with or without *S. aureus* for 4 h. Results are normalized to non-treated wild-type animals. Validation of RNA-seq using three biological replicates per condition and three technical replicates per biological repeat. Significance was determined using one-way ANOVA with the application of the Bonferroni correction (^*^*p*<0.05, ^**^*p*<0.01, \*\*\**p*<0.001, ^****^*p*<0.001).

### AMPK regulates TFEB/TFE3-mediated innate immune response

Based on our previous and current data, we tested whether the transcriptional innate immune response observed upon loss of FLCN is similar to a gain in AMPK activity in mammalian cells using GSK-621, a specific AMPK activator [38, 39]. In MEFs, we show that GSK-621 activated AMPK, as shown by increased downstream target p-ACC, without inhibiting mTOR signaling as measured by immunoblotting for the phosphorylated forms of p70S6K and 4EBP1 (Figure 6A). Such activation was accompanied by a significant increase in the nuclear translocation of TFEB and TFE3 (Figures 6B-C), which was lost in AMPKα1/α2 double knock out (DKO) MEFs (Figures 6B-C), confirming the specific activation of AMPK by GSK-621. Additionally, we show that IL-6, a TFEB/TFE3 target, was transcriptionally upregulated when treated with GSK-621 and its expression was abrogated upon down-regulation of TFEB/TFE3 using shTFEB/TFE3 (Figure 6D) implying that AMPK impinges on TFEB/TFE3-mediated transcription in mammalian cells similarly to what we have observed in *C. elegans*. Moreover, treatment of RAW264.7 macrophages with GSK-621 activated AMPK without affecting mTOR signaling (Figure 6E), promoted the nuclear translocation of TFEB (Figure 6F), and led to a strong increase in production and secretion of various cytokines and chemokines even in the absence of LPS treatment or pathogen infection (Figure 6G). To substantiate our findings in a more physiological context, we tested whether acute LPS treatment of macrophages could affect cellular bioenergetics, which could be sensed by AMPK. Indeed, we observed an acute reduction in cellular ATP levels (Figure 6H), accompanied by AMPK activation (Figure 6I), and a significant increase in TFEB nuclear localization (Figure 6J) as early as 30 minutes after addition of LPS in RAW macrophages.

**Fig 6:**
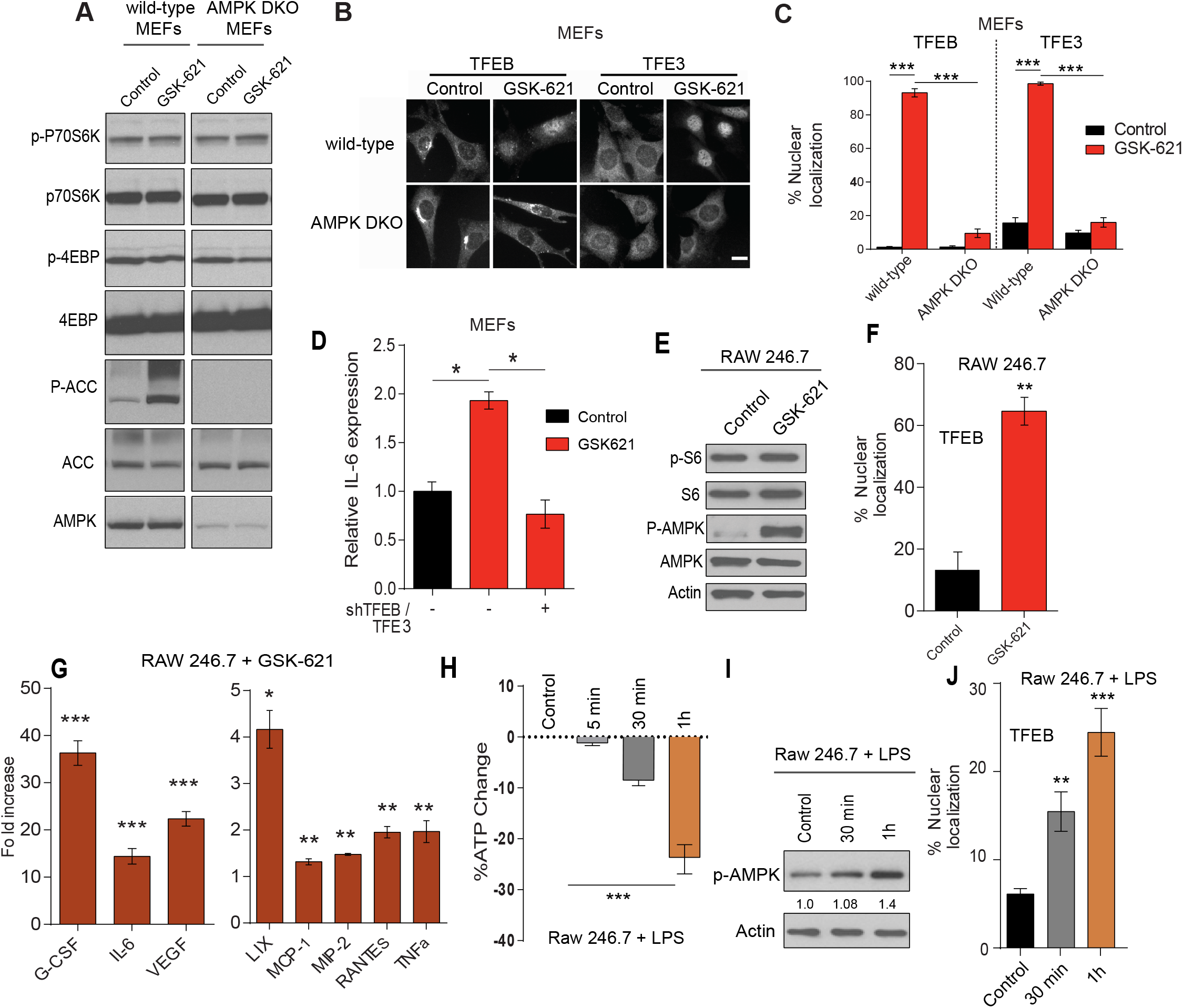
AMPK regulates TFEB/TFE3-mediated innate immune response. (A) Immunoblot of wild-type or AMPKα1/α2 double knock out (DKO) MEFs stimulated with the AMPK activator; GSK-621 (30 μM) for 1 h. (B) Representative images of TFEB and TFE3 staining in wild-type and AMPK DKO MEFs before and after treatment with GSK-621 (30 μM) for 1 h. Scale bars represent 20 μm. (C) Quantification of the percentage of cells showing TFEB and TFE3 nuclear staining of the conditions described in (B). (D) Relative IL-6 mRNA levels measured by qRT-PCR in wild-type MEFs transfected with EV or shTFEB/TFE3, stimulated with GSK-621 for 2 h. Data represents the average of three independent experiments, each done in triplicates ± SEM. Significance was determined using one-way ANOVA with the application of Bonferroni correction (^*^*p*<0.05, ^***^*p*<0.001). (E) Immunoblot analysis of RAW264.7 macrophages treated with GSK-621 (30 μM) for 2 h. (F) Quantification of the percentage of RAW264.7 cells showing TFEB nuclear staining of the conditions described in (D). (G) Quantification of the significant fold increases in cytokine and chemokine protein levels in RAW264.7 macrophages, treated with GSK-621 (30 μM) for 2 h as compared to control. (H) Fold change in ATP levels in RAW264.7 treated with LPS (1 μg/ml) for up to 1 h as measured by CellTiter-Glo Luminescent Cell Viability Assay. (I) Immunoblot analysis of RAW264.7 macrophages treated with LPS (1 μg/ml) for up to 1 h. (J) Quantification of the percentage of RAW264.7 cells with TFEB nuclear localization of the conditions described in (H). Data represent the average of three independent experiments, each done in triplicates ± SEM. Significance was determined using student’s t-test (^*^*p*<0.05, ^**^*p*<0.01, ^***^*p*<0.001).

Collectively, both the mammalian and worm results demonstrate an important role for AMPK in the regulation of the innate host immune response through TFEB/TFE3 activation.

## Discussion

We have previously shown that loss of FLCN activates AMPK, increasing the resistance to oxidative stress, heat, anoxia, hyperosmotic stresses, and obesity in *C. elegans* and mammalian models [28-31, 40]. Here, we report for the first time evidence supporting a novel and evolutionary conserved role of FLCN in innate host defense mediated through AMPK and TFEB/TFE3 activation.

Given that the gene profile upon osmotic stress mimics that of pathogen infection [36], we found a significant overlap in the transcriptional profile in *flcn-1* mutant animals when compared to wild-type animals infected with pathogens. We report that *flcn-1* mutant animals confer resistance to pathogen infection through nuclear localization and activation of HLH-30. Increased nuclear localization and activation of TFE3 were previously reported in renal tumors from Birt Hogg-Dubé syndrome patients, a syndrome associated with a germline mutation of the FLCN gene [41]. Subsequent studies further supported a role for FLCN in the cytoplasmic retention of TFE3 and TFEB [42-45]. The mechanisms through which TFEB and TFE3 are regulated in response to nutrient status have been characterized. Most studies to date suggested that mTORC1-dependent phosphorylation of TFEB causes cytoplasmic retention of this transcription factor under nutrient-rich conditions. Inhibition of mTORC1 activity upon nutrient starvation has been associated with hypo-phosphorylated forms and nuclear accumulation of TFEB and TFE3 inducing the up-regulation of genes involved in autophagy and lysosomal biogenesis, and thus favoring cell survival and adaption to stress [14, 16, 17, 43, 44, 46]. The link between FLCN and mTOR has been previously proposed, where FLCN was identified as a GTP-Activating Protein (GAP) for Ras-related GTPase (Rag)C/D, and a Guanine Exchange Factor (GEF) for RagA/B, which ultimately activates mTOR [44, 47]. The yeast ortholog of FLCN, Lst7, also acts as a GAP for yeast RagC/D ortholog Gtr2 [48]. Conversely, FLCN-deficient tumors were shown to exhibit activated mTOR while acute loss of FLCN in cellular systems led to mTOR inhibition [22, 44, 47-49], suggesting that FLCN’s role in this process is cell and context-dependent, and might vary in response to different environmental signals.

Our current work reveals that pathogen-induced regulation of TFEB and TFE3 activation appears to have different dynamics than that of starvation-induced regulation [17]. Using both *C. elegans* and mammalian models, we show that the FLCN/AMPK axis and the mTOR axis impinge differently and independently on TFEB and TFE3 activation status. We show in nematodes that AMPK regulated the nuclear localization of HLH-30 and the transcription of anti-microbial genes. In mammalian cells, we show that AMPK activation led to the transcriptional up-regulation of pro-inflammatory cytokines through the nuclear translocation and activation of TFEB/TFE3. AMPK has been shown to govern lineage specification by promoting autophagy and lysosomal biogenesis through transcriptional mechanisms including TFEB [50]. Although no direct link between AMPK and TFEB has been reported, AMPK was thought to activate TFEB through inhibition of mTORC1 [50]. Conversely, and in support of our observed results herein, it has been shown that while starvation-mediated activation of TFEB/TFE3 involved mTORC1, their pathogen-induced activation appeared to be mTORC1 independent [18].

Bacterivorous nematodes, such as *C. elegans* induce the expression of transcriptional host-defense responses including the HLH-30/TFEB pathway that promote organismal survival [6, 37, 51-53]. However, these invertebrates appear to lack the NLR and TLR pathogen sensing pathways as well as NF-κB and other transcription factor pathways that regulate innate immunity in higher organisms [3, 54]. Our findings shed light on an ancient, highly conserved pathogen sensing and signal transduction mechanism, which involves AMPK and the transcription factor TFEB/TFE3. LPS, which is part of the outer membrane of Gram-negative bacteria, was shown to inhibit respiration and energy production in cells and isolated mitochondria [55-58]. We show here, that LPS treatment of macrophages leads to an acute reduction of cellular energy levels resulting in AMPK activation, induction of TFEB/TFE3 and inflammatory cytokines (Figure 6H-K; Figure 3H, I). Therefore, this ancient pathogen-sensing pathway may have evolved by the fact that pathogen infection leads to an energy shortage, which is sensed by cellular AMPK. Activated AMPK will in turn promote TFEB/TFE3 nuclear translocation and induction of an innate host defense.

It remains to be elucidated how exactly loss of FLCN leads to AMPK activation on a mechanistic level. FLCN is a GAP for RagC/D, which ultimately activates mTOR [44, 47]. Loss of FLCN leads to permanent mTOR inhibition with respect to activation of lysosomal biogenesis and autophagy via TFEB/TFE3, whereas mTOR-mediated signaling towards protein synthesis appears not to be affected [45]. We showed previously that loss of binding of FLCN to FNIP/AMPK via introduction of a phospho-mutant of FLCN as well as knockdown or complete loss of FLCN leads to permanent activation of AMPK with respect to autophagy via ULK-1, mitochondrial biogenesis via PGC-1α, glycolysis and angiogenesis via HIF-1α, glycogen metabolism upon osmotic stress, and resistance to obesity via induction of functional beige adipose tissue [28, 29, 31, 40]. However, more detailed work needs to be performed to fully understand the role of FLCN binding to FNIP and AMPK and its regulation in the role of AMPK activation.

How AMPK activates TFEB and TFE3 upon depletion of FLCN remains unknown. However, it is likely that AMPK activation under pathogen-induced conditions regulates TFEB and TFE3 activation distinctly from the mTOR pathway. Recent studies have shown that the kinases PLC-1 and DKF-1, the *C. elegans* orthologs of mammalian PLC and PRKD1/PKD, respectively, are required for HLH-30 activation during infection of nematodes with *S. aureus* [21]. A similar mechanism of TFEB activation in mouse macrophages infected with pathogens involved the PRKCA/PKCa axis demonstrating that TFEB activation in response to pathogen infection is conserved throughout evolution [21]. It appears that TFEB/TFE3 are controlled by a panel of kinases and phosphatases that depending on the environmental cues exhibit different downstream responses. Quantitative proteomics have identified over 20 phosphorylation sites on TFEB and TFE3, and although not directly tested in this current study, assessing their direct phosphorylation by AMPK could provide information about the contribution of AMPK in TFEB regulation.

The mechanisms through which TFEB and TFE3 confer pathogen resistance are still being deciphered. While TFEB/TFE3 activation were previously reported not to affect pathogen burden over the course of infection, they appear to regulate the mechanisms of tolerance to infection through autophagy/lysosomal pathways that enhance ability of the host to survive upon pathogen invasion [6]. Moreover, induction of lysosomal pathways have been demonstrated to enhance the phagocytic capacity of innate immune effector cells [59]. In this study, we show that down-regulation of FLCN in murine macrophages enhances their phagocytic activity and prompts a metabolic transformation toward increased cellular bioenergetics, which might further enhance the innate immune response. FLCN/AMPK-mediated increase in autophagic flux and AMPK/TFEB-mediated increase in lysosomal biogenesis are likely to contribute to metabolic fitness of infected cells and increased phagocytosis in macrophages. Interestingly, and in line with our results, it was recently proposed that the activation of the Fcγ receptor in macrophages enhances lysosome-based proteolysis and killing of phagocytosed *E. coli* and this activation induces the nuclear translocation of TFEB accompanied by an increase the expression of specific lysosomal proteins. Notably, TFEB silencing represses the Fcγ-receptor-mediated enhancements in degradation and bacterial killing [60]. Hence, further studies are required to elucidate precisely how TFEB/TFE3 regulation through FLCN/AMPK axis affects host tolerance of infection in nematodes and in mammals.

Patients affected with BHD syndrome are at risk of developing bilateral, multifocal renal tumors, skin tumors and lung cysts [61]. In addition, chromosomal translocations leading to TFE3 or TFEB over-activation were reported in sporadic juvenile and advanced renal cell carcinoma (RCC) [62]. Hence, it is tempting to speculate whether loss of FLCN and AMPK activation in humans induce a chronic inflammatory response and thereby promoting cancer progression, similar to reported cancer cases where innate immune response pathway such as NF-κB is over-activated [63].

Furthermore, in this study we place AMPK at the center of FLCN-TFEB/TFE3 axis. Several direct AMPK activators are being developed for treatment of type-2 diabetes, obesity, and metabolic syndrome [64]. We propose that some of these specific AMPK activating compounds could be repurposed to enhance host defense against pathogens or treat other immunodeficiency syndromes through AMPK-mediated activation of TFEB/TFE3, providing new druggable strategies in innate immune modulation and therapy of bacterial infections. To this end, pharmacological inhibition of mTOR is currently being investigated in human clinical trials to treat age-associated immune dysfunction, also dubbed “immune senescence” (resTORbio, Inc).

## Acknowledgements

We thank Audrey Kapelanski-Lamoureux for technical support, Javier Irazoqui for kindly providing the *hlh-30(tm1978)* mutant strain, Malene Hansen for the *hlh-30*::GFP transgenic line, Joaquin Madrenas and Eric Deziel for providing us the MW2 *S. aureus* and PA14 *P. aeruginosa* bacterial strains, respectively. We thank Dr. Yong Chen and Dr. Marjan Gucek (Proteomic Core Facility, NHLBI, NIH) for their assistance with mass spectrometry analysis. We thank Nahum Sonenberg and Marie-Claude Gingras for the critical reading of the manuscript. We acknowledge the *Caenorhabditis* Genetic Center for *C. elegans* strains. E.P., L.E., M.P., and T. V. were supported by Rolande and Marcel Gosselin studentship and the MICRTP training grant, Dr. Victor K.S. Lui Fellowship and Dr. Michael D’Avirro from GCRC, FRQS, and Canderel studentship award, respectively. This work was supported by grants to A.P. from Myrovlytis Trust, Kidney Foundation of Canada and Terry Fox Research Institute. The Goodman Cancer Research Centre Metabolomics Core Facility is supported by the Canada Foundation of Innovation and Terry Fox Research Institute. R.J. and E.M. were supported by CIHR grant MOP-142259. J.A.M. and R.P. were supported by the Intramural Research Program of the National Institutes of Health, National Heart, Lung, and Blood Institute (NHLBI).

## Material and methods

### *C. elegans* strains, maintenance, and RNAi treatments

All strains used in this study are described in supplementary table S20 (Table S20). Nematodes were maintained and synchronized using standard culture methods [65]. The RNAi feeding experiments were performed as described in [66], and bacteria transformed with empty vector were used as control. For all RNAi experiments, phenotypes were scored with the F1 generation.

### Pathogen resistance assay

To measure pathogen stress resistance, synchronized L4 worms were transferred to Tryptic Soya Agar (TSA) plates with 8 μg/ml Nalidixic acid that were seeded with 1:50 *S. aureus* MW2 bacteria incubated at 37°C for 3 h [67]. Survival was measured daily by transferring worms onto new plates. To measure stress resistance to *P. aeruginosa* PA14, synchronized L4 worms were transferred to Slow Killing (SK) plates [67]. Worms were transferred to new plates every 24 h to monitor survival. Worms that responded with movement upon being touched by a platinum wire were considered alive. Assays were performed in triplicate plates per condition with 30 animals per plate, in three independent experiments.

### HLH-30 nuclear translocation assay

The *hlh-30p::hlh-30::GFP* was kindly provided by Malene Hansen’s Lab. The *flcn-1(ok975); hlh-30p::hlh-30*::GFP, *aak-2(gt-33)*; *hlh-30p::hlh-30*::GFP and *aak-1(tm1944);aak-2(gt-33); hlh-30p::hlh-30*::GFP strains, respectively, were obtained using standard genetic crossing strategies. 30-40 worm eggs were transferred to 35 mm regular NGM plates seeded with OP50. Synchronized young adult animals expressing the HLH-30:GFP transgene were transferred to TSA plates seeded with 1:20 *S. aureus* MW2 bacteria on the day of the experiment. Worms displaying HLH-30 translocation were scored. For the *let-363* RNAi based experiments, synchronized animals were grown on RNAi plates and were used at the young adult stage for infection with *S. aureus* MW2 bacteria. Translocation was counted using a fluorescent dissecting microscope at indicated timepoints and imaged using Zeiss confocal laser scanning microscope. Images were taken within the first 5 min because mounting stress also induces HLH-30 nuclear translocation.

### RNA extraction and real-time PCR in *C. elegans*

Synchronized young adult nematodes were exposed to pathogenic *S. aureus* bacteria or OP50 seeded plates for 4 h, harvested and washed with M9 buffer. Pellets were flash frozen in liquid nitrogen. Total RNA was extracted with Trizol. iScript Supermix from Bio-Rad was used to reverse transcribe 1 μg RNA samples. Quantitative RT-PCRs were performed as previously described with small modifications [28]. Bio-Rad SYBR Green mix was used and qPCRs were performed on the Roche LightCycler 480 machine. Three housekeeping genes were used to confirm changes in gene expression, *cdc-42, pmp-3* and Y45FD10.4 [68].

### Microarray experiment and gene overlap analysis

Synchronized L4/young adult wild-type and *flcn-1(ok975)* animals were harvested and RNA was extracted using Trizol and purified on Qiagen RNeasy columns. Total RNA samples were then hybridized onto Agilent gene chips. Fold change values were calculated using the mean of both data sets. Agilent files were uploaded into the FlexArray software at Genome Quebec for analysis. Three replicates were normalized and analyzed for each condition. Fold change was determined and p-value was obtained using a standard student’s t-test. Differentially expressed genes were compared to other studies; hyperosmotic stress [35] and pathogen infection [36, 37] using the “compare two lists” online tool at http://www.nemates.org/MA/progs/Compare.html. The significance of the overlap and enrichment scores was determined via hypergeometric distribution method using http://nemates.org/MA/progs/overlap_stats.html. The number of genes in the *C. elegans* genome was considered 19,735.

### RNA seq method

Synchronized wild-type and *aak-1(tm1944); aak-2(ok524)* animals were harvested at the late L4 stage, washed with M9, and flash frozen in liquid nitrogen. RNA was extracted using Trizol and purified using Qiagen RNeasy columns. RNA samples were processed for RNA-seq analysis at Novogene Inc.

### Reagents, chemicals, and antibodies

LPS derived from Escherichia Coli endotoxin (0111:B4, InvivoGen, San Diego, CA, USA) was dissolved in PBS (5 mg/ml) by sonication for 2 min, aliquoted and stored at −80°C until use. All LPS preparations were free of protein or lipoprotein contaminants. LPS was used at a final concentration of 1 μg/ml. Recombinant mouse TNF-α was obtained from Biolegend (#575206) with a stock concentration of 0.2 mg/ml and was used at a final concentration of 10 ng/ml dissolved in 10% DMEM. GSK-621 was obtained from APExBIO or Selleckchem (Houston, Texas, USA) and dissolved in DMSO to a stock concentration of 30 mM and used at a final concentration of 30 μM for MEFs. Torin1 was obtained from Tocris Bioscience (Bristol, UK) and dissolved in DMSO to a stock concentration of 1 mM and used at a final concentration of 1 μM. The final DMSO concentration never exceeded 0.1% and this concentration was shown to have no detrimental effect on all the studied cells.

The mouse FLCN polyclonal antibody was generated by the McGill animal resource center services through injecting purified GST-FLCN recombinant protein in rabbits. ß-Actin (sc-47778; Santa Cruz Biotechnology), Tubulin (T9026; Sigma), LaminA (SC-71481; Santa Cruz Biotechnology), AMPKα (2532; Cell Signaling Technology), p-AMPKα (Thr172) (2531; Cell Signaling Technology), ACC (3676; Cell Signaling Technology), p-ACC (S79) (3661; Cell Signaling Technology), TFEB (A303-673A; Bethyl Laboratories), TFE3 (14779S; Cell Signaling Technology and HPA023881; Sigma-Aldrich), p70S6K (2708; Cell Signaling Technology), p-p70S6K (9205; Cell Signaling Technology), S6 (2217; Cell Signaling Technology), p-S6 (4858; Cell Signaling Technology), 4EBP1 (9644; Cell Signaling Technology), p-4EBP1 (9456; Cell Signaling Technology) antibodies are commercially available.

### Cell culture

Primary MEFs were isolated from C57BL/6 E12.5 *Flcn* floxed mice (generously provided by Dr. Laura S. Schmidt, NCI, Bethesda, MD, USA) and cultured as described in [29]. Flcn wild-type and knockout (KO) MEFs were generated after immortalization of primary Flcn Flox/Flox MEFs with retroviral infection of SV40 large T (hygromycin B) and retroviral infection of CD8 or CD8-Cre recombinase, followed by FACS sorting of CD8 positive cells. AMPK DKO MEFs and their wild-type counterpart cells were generously provided by Dr. Benoit Viollet (Institut Cochin INSERM, Paris, France). RAW 264.7 cells (termed RAW cells), a murine macrophage cell line (ATCC CRL-24) were generously provided by Dr. C. Krawczyk (McGill University, Montreal, Canada). RAW 264.7 TFEB/TFE3 DKO cells and their EV counterpart cells were generously provided by Dr. Rosa Puertollano (National Institutes of Health, Bethesda, MD, USA). Cell lines were maintained in Dulbecco’s modified Eagle’s medium (DMEM) supplemented with 10% fetal bovine serum (FBS) (Wisent), 100 U/ml penicillin and 100 μg/ml streptomycin (Invitrogen) in 5% CO_2_ at 37°C. MEFs and RAW cells were stably downregulated for TFEB/TFE3 or FLCN, respectively, using the Mission lentivirus shRNA empty vector (shEV), shTFEB (TRCN0000013110; Sigma-Aldrich), shTFE3 (TRCN0000232151; Sigma-Aldrich), or shFLCN (TRCN0000301434; Sigma-Aldrich).

### Quantitative real-time RT-PCR in mammalian cells

FLCN wild-type, FLCN KO MEFs, and RAW 264.7 cells were seeded in triplicates in 6- well plates at 3 × 10^5^ cells per well in DMEM medium supplemented with 10% FBS. After incubation for 24 h at 37°C, 5% CO_2_, cells were treated with TNF-α, LPS, GSK-621 or vehicle for 2, 3, or 24 h. Cells were then collected, and total RNA was isolated and purified using Total RNA Mini Kit (Geneaid) according to the manufacturer’s instructions. For quantitative real-time PCR analysis, 1 μg of total RNA was reverse-transcribed using the SuperScript III kit (Invitrogen). SYBR Green reactions using the SYBR Green qPCR supermix (Invitrogen) and specific primers (table below) were performed using an AriaMX Real-time PCR system (Agilent Technologies). Relative expression of mRNAs was determined after normalization against housekeeping gene RPLP0 or B2M.

### Mouse Protein Cytokine Array

RAW 264.7 cells were seeded in triplicates in 6-well plates at 1 × 10^6^ cells per well in DMEM medium supplemented with 10% FBS. After incubation for 24 h at 37 °C, 5% CO_2_, cells were treated with LPS or GSK-621 or vehicle for 3, or 24 h, and the conditioned medium was harvested and centrifuged at 1,500 × *g* to remove cell debris. 32 cytokine/chemokine/growth factor biomarkers were simultaneously quantified by using a Discovery Assay^®^ called the Mouse Cytokine Array/Chemokine Array 32-Plex (Eve Technologies Corp, Calgary, AB, Canada). The multiplex assay was performed by using the Bio-Plex™ 200 system (Bio-Rad Laboratories, Inc., Hercules, CA, USA), and a Milliplex Mouse Cytokine/Chemokine kit (Millipore, St. Charles, MO, USA) according to the manufacturers protocol. The 32-plex consisted of Eotaxin, G-CSF, GM-CSF, IFNγ, IL-1α, IL-1β, IL-2, IL-3, IL-4, IL-5, IL-6, IL-7, IL-9, IL-10, IL-12 (p40), IL-12 (p70), IL-13, IL-15, IL-17, IP-10, KC, LIF, LIX, MCP-1, M-CSF, MIG, MIP-1α, MIP-1β, MIP-2, RANTES, VEGF. The change in the cytokine levels in FLCN KO medium was normalized against their respective wild-type medium.

### Protein extraction and immunoblotting

For AMPK immunoblotting, cells were washed twice with cold PBS, lysed in AMPK lysis buffer (10 mM Tris-HCl (pH 8.0), 0.5 mM CHAPS, 1.5 mM MgCl_2_ 1 mM EGTA, 10% glycerol, 5 mM NaF, 0.1 mM Na_3_VO_4_, 1 mM benzamidine, 5 mM NaPPi), supplemented with complete protease inhibitor (Roche) and DTT (1 mM), and cell lysates were cleared by centrifugation at 13000 × *g*. For all other immunoblotting, cells were washed twice with cold PBS and lysed directly in Laemmli buffer (62.5 mM Tris-HCl (pH 6.8), 2% (w/v) sodium dodecyl sulphate, 10% (v/v) glycerol, 5% (v/v) β-mercaptoethanol and 0.01% (w/v) bromophenol blue). Proteins were separated on SDS-PAGE gels and revealed by western blot as we previously described [29] using the antibodies listed above.

### Subcellular fractionation

Cells were washed and resuspended in PBS and pelleted 5 min at 300 × *g* at 4°C. The pellet was lysed in NP-40 lysis buffer (50 mM Tris-HCl (pH 7.4), 0.5 % NP-40, 5 mM MgCl_2_ 140 mM KCl, 5 mM NaF, 0.1 mM Na_3_VO_4_, 1 mM benzamidine, 5 mM NaPPi, 1x protease inhibitors) and pelleted 5 min at 1000 × *g*. Supernatant was collected as the cytosolic fraction. The pellet was lysed in nuclear lysis buffer (25 mM Tris-HCl (pH 7.4), 0.5 % Triton X-100, 0.5% SDS, 5 mM MgCl_2_ 140 mM KCl, 5 mM NaF, 0.1 mM Na_3_VO_4_, 1 mM benzamidine, 5 mM NaPPi, 1 × protease inhibitors), sonicated 15 min with 30 sec bursts and collected as the nuclear + membrane fraction.

### Immunofluorescence

Cells were washed with PBS and fixed in petri dishes with 3.7% formaldehyde at room temperature for 30 min. After fixation, cells were washed twice with PBS and then permeabilized with 0.3% Triton X-100 in PBS at room temperature for 5 min. Cells were incubated in 10% BSA in PBS for 1 h and then with TFEB or TFE3 primary antibody in 1.5% BSA in PBS for 2 h at 37°C. Cells were washed three times with PBS and incubated with the corresponding secondary antibodies conjugated to Alexa Fluor 488 in 1.5% BSA in PBS for 30 min at 37°C. Cells were washed three times with PBS and incubated with DAPI (0.1 μg/ml) in PBS for 15 min at room temperature. PBS-washed dishes were covered with cover slips and observed with Axioskop microscope (Zeiss).

### Metabolic Assays

Glucose production and lactate consumption was measured using a NOVA Bioanalysis flux analyzer or the Eton Bioscience kit (Eton Bioscience, Charlestown, MA, USA) according to the manufacturer’s instructions. OCR and the ECAR of cells were measured using an XF96 Extracellular Flux Analyzer (Seahorse Bioscience, Boston, MA, USA) as previously described [69]. In brief, EV or shFLCN RAW264.7 were plated at 100,000 cells/well in growth medium for 24 h. After 24 h, cells were incubated in non-buffered DMEM containing 25 mM glucose and 2 mM glutamine in a CO2-free incubator at 37°C for 2 h to allow for temperature and pH equilibration before loading into the XF96 apparatus. XF assays consisted of sequential mix (3 min), pause (3 min), and measurement (5 min) cycles, allowing for determination of OCR/ECAR every 10 min. After establishing a baseline, Oligomycin (10 uM), FCCP (15 μM), and Rotenone/Antimycin A (1 μM, and 10 μM, respectively) were added sequentially.

### ATP quantification

Cells were plated in triplicates in 96-well plates. After 24 h, cells were lysed and mixed for 10 min according to manufacturer’s instructions (CellTiter-Glo luminescent cell viability assay, Promaga). Luminescence was measured using Fluostar Omage (BMG Labtech) directly in plates.

### Phagocytosis Assay

Phagocytosis in EV or shFLCN RAW264.7 cells was assessed using Red pHrodo *S.aureus* BioParticles conjugate assay (Thermofisher) according to the manufacturer’s protocol. In brief, EV or shFLCN RAW264.7 were plated in triplicates in 96-well plates at 80,000 cells/well in growth medium for 2 h before treatment. After 2 h, cells were treated with the BioParticles (after homogenization in serum-free DMEM) at a final concentration of 1 mg/ml and incubated at 37 °C for 3 h. Subsequently, cells were collected and analyzed using FACSDiva analyzer (Becton Dickison).

### Statistical Analyses

Data are expressed as mean ± SEM. Statistical analyses for all data were performed using student’s t-test for comparisons between 2 groups, one-way ANOVA for comparisons between 3 or more groups, and Log-rank Mantel Cox test for survival plots, using GraphPad Prism 7 software. Statistical significance is indicated in figures (^*^*p*<0.05, ^**^*p*<0.01, ^***^*p*<0.001, ^****^*p*<0.0001) or included in the supplemental tables.

---

**Fig S1:**
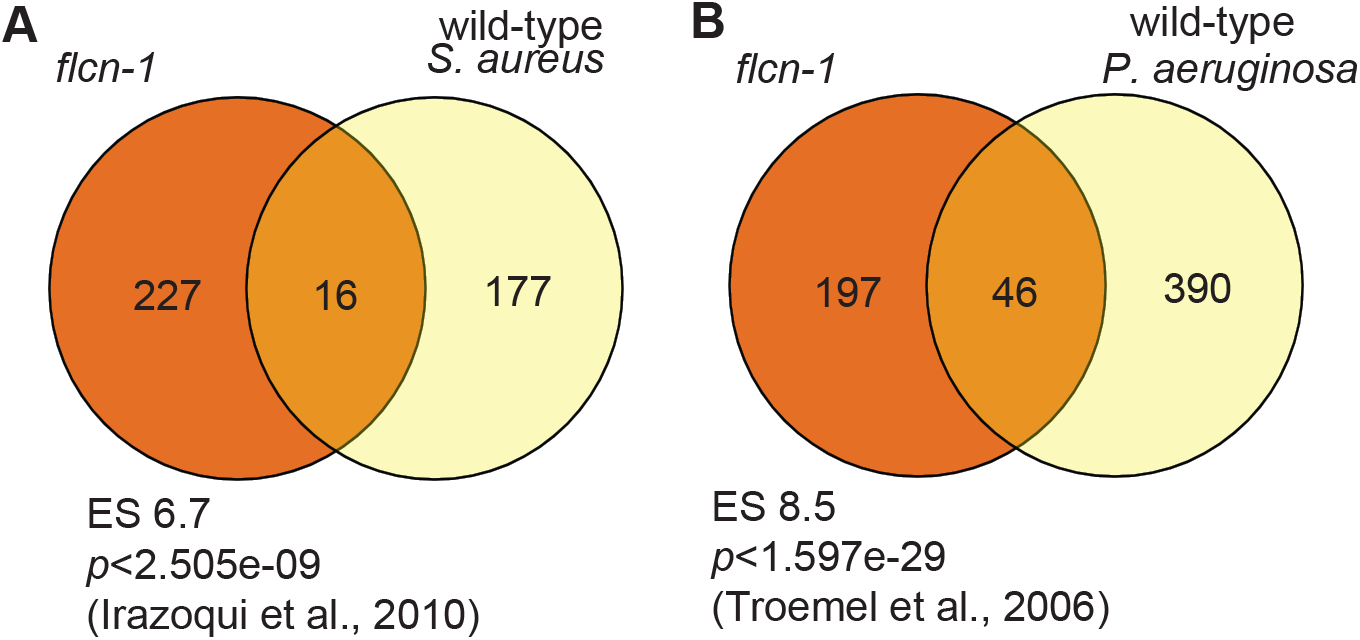
Transcriptional profile of *flcn-1* prior to stress overlaps with profiles of wild-type animals infected with pathogens and correlates with a pathogen resistance phenotype. (A, B) Venn diagrams showing the overlap of genes upregulated in *flcn-1(ok975)* animals at basal level and genes upregulated in wild-type animals following treatment with *S. aureus* from [36] (A) or *P. aeruginosa* from [37] (B). Comparisons were performed using the “compare two lists” online software and the significance and ES (enrichment scores) were obtained using “nemates” software.

**Fig S2:**
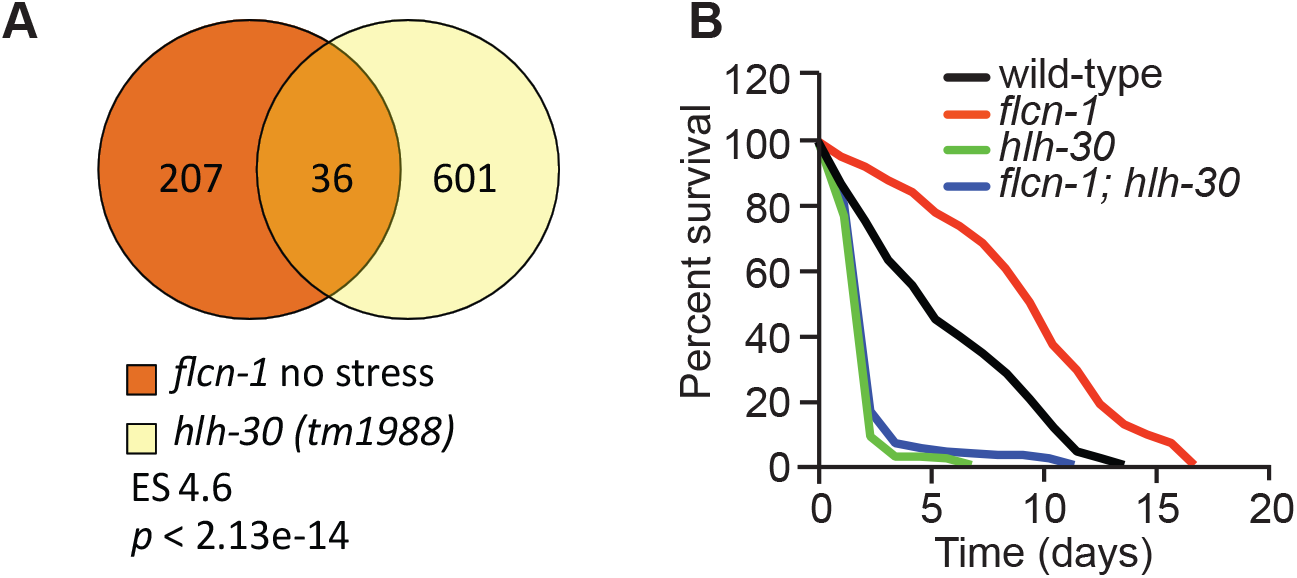
Role of *hlh-30* downstream of *flcn-1*. (A) Venn diagrams showing the overlap of genes upregulated in *flcn-1(ok975)* animals at basal level and downregulated in *hlh-30 (tm1978)* mutant nematodes. Comparisons were performed using the “compare two lists” online software and the significance and ES (enrichment scores) were obtained using “nemates” software. (B) Percent survival of indicated strains to 400 mM NaCl stress. Refer to Table S9 for details on number of animals utilized and number of repeats Statistics obtained by Mantel-Cox analysis on the pooled curve.

**Fig S3:**
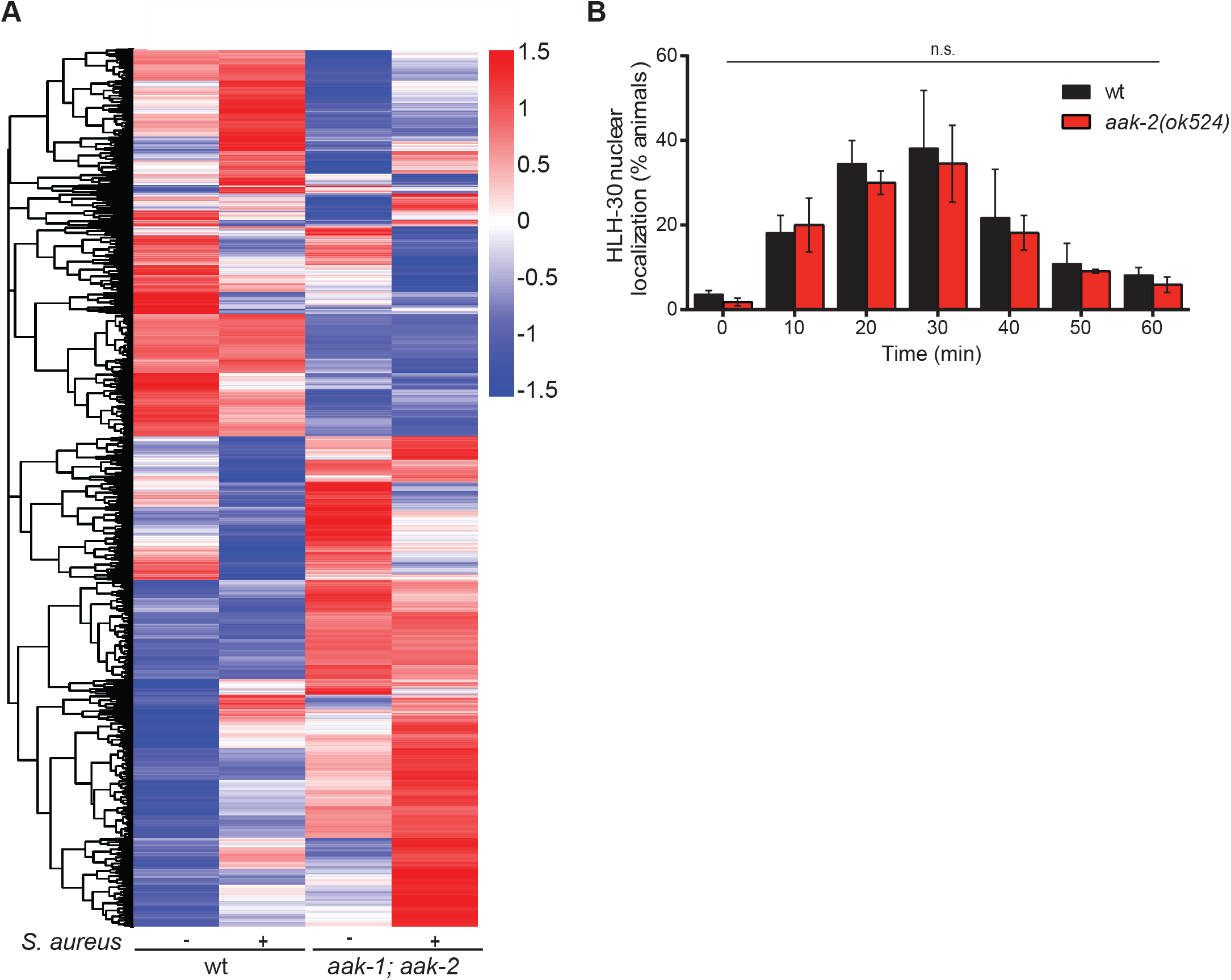
RNA Seq heat map and gene ontology analysis in wild-type and *aak-1(tm1944); aak-2(ok524)* at basal level and upon *S. aureus* infection. (A) Heat map showing differential gene expression in wild-type and *aak-1(tm1944); aak-2(ok524)* mutant animals grown on OP50 or exposed to *S. aureus* for 4 h. Red color indicates genes that are differentially upregulated while blue color indicates gene sets that are downregulated in comparison to untreated wild-type animals. RNAseq data were analysed by Novogene Inc. using DEGseq 1.12.0 (B) Nuclear translocation of HLH-30 in *aak-2(ok524); hlh-30::GFP* at basal level and upon *S. aureus* infection. Data represent the mean ± SEM with 3 independent repeats, n ≥ 30 animals/condition for every repeat. Significance was determined using student’s t-test.

**Table S1: List of genes upregulated in *flcn-1(ok975)* mutant animals in comparison to wild-type at basal level.**

**Table S2: GO analysis of genes upregulated in *flcn-1(ok975)* mutant animals at basal level.**

**Table S3: List of genes with known antimicrobial and defense functions upregulated in *flcn-1(ok975)* at basal level; selection based on GO annotations.**

**Table S4: List of genes downregulated in *flcn-1(ok975)* mutant animals at basal level.**

**Table S5: GO analysis of genes downregulated in *flcn-1(ok975)* mutant animals at basal level.**

**Table S6: Overlapping genes upregulated in *flcn-1(ok975)* animals and animals infected with *S. aureus*.**

**Table S7: Overlapping genes upregulated in *flcn-1(ok975)* animals and animals infected with *P. aeruginosa.***

**Table S8: (A) Mean survival on *S. aureus*: results and statistical analysis. (B) Mean survival on *P. aeruginosa*: results and statistical analysis.**

**Table S9: Mean survival on 400mM NaCl: results and statistical analysis**

**Table S10: List of overlapping genes upregulated in *flcn-1(ok975)* mutant animals at basal level and *S. aureus* hlh-30-dependent genes.**

**Table S11: Genes upregulated in wild-type animals treated with *S. aureus* for 4 h in comparison to animals grown on OP50 *E. Coli.***

**Table S12: List of genes downregulated in wild-type animals treated with *S. aureus* for 4 h in comparison to animals grown on OP50.**

**Table S13: List of genes upregulated in *aak-1(tm1944); aak-2(ok524)* animals in comparison to wild-type animals at basal level.**

**Table S14: List of genes downregulated in *aak-1(tm1944); aak-2(ok524)* animals in comparison to wild-type animals at basal level.**

**Table S15: List of genes downregulated in *aak-1(tm1944); aak-2(ok524)* animals treated with *S. aureus* in comparison to wild-type animals treated with *S. aureus*.**

**Table S16: AMPK-dependent genes determined by overlap between genes upregulated in wild-type animals upon *S. aureus* infection and downregulated in infected *aak-1(tm1944); aak-2(ok524)* mutant animals.**

**Table S17: GO analysis of AMPK-dependent genes obtained by overlap between genes upregulated in wild-type animals upon S*. aureus* infection and downregulated in infected *aak-1(tm1944); aak-2(ok524)* mutant animals. This table includes a histogram of functional gene ontology analysis of genes induced by *S. aureus* in wild-type animals and downregulated in *aak-1(tm1944); aak-2(ok524)* mutant animals upon infection.**

**Table S18: Genes downregulated in *aak-1(tm1944); aak-2(ok524)* and *hlh-3(tm1978)* mutant animals upon *S. aureus* infection in comparison to wild-type.**

**Table S19: GO analysis of the overlap between infection genes regulated by AMPK and HLH-30.**

**Table S20: List of *C. elegans* strains used in this study.**

## References

1. Akira, S., S. Uematsu, and O. Takeuchi, Pathogen recognition and innate immunity. Cell, 2006. 124(4): p. 783–801.

2. Hoffmann, J.A., The immune response of Drosophila. Nature, 2003. 426(6962): p. 33–8.

3. Irazoqui, J.E., J.M. Urbach, and F.M. Ausubel, Evolution of host innate defence: insights from Caenorhabditis elegans and primitive invertebrates. Nat Rev Immunol, 2010. 10(1): p. 47–58.

4. Medzhitov, R., Recognition of microorganisms and activation of the immune response. Nature, 2007. 449(7164): p. 819–26.

5. Medzhitov, R., Damage control in host-pathogen interactions. Proc Natl Acad Sci U S A, 2009. 106(37): p. 15525–6.

6. Visvikis, O., et al., Innate host defense requires TFEB-mediated transcription of cytoprotective and antimicrobial genes. Immunity, 2014. 40(6): p. 896–909.

7. Lapierre, L.R., et al., The TFEB orthologue HLH-30 regulates autophagy and modulates longevity in Caenorhabditis elegans. Nat Commun, 2013. 4: p. 2267.

8. Rehli, M., et al., Cloning and characterization of the murine genes for bHLH-ZIP transcription factors TFEC and TFEB reveal a common gene organization for all MiT subfamily members. Genomics, 1999. 56(1): p. 111–20.

9. David, R., Autophagy: TFEB perfects multitasking. Nat Rev Mol Cell Biol, 2011. 12(7): p. 404.

10. Raben, N. and R. Puertollano, TFEB and TFE3: Linking Lysosomes to Cellular Adaptation to Stress. Annu Rev Cell Dev Biol, 2016. 32: p. 255–278.

11. Sardiello, M., Transcription factor EB: from master coordinator of lysosomal pathways to candidate therapeutic target in degenerative storage diseases. Ann N Y Acad Sci, 2016. 1371(1): p. 3–14.

12. Settembre, C., et al., TFEB controls cellular lipid metabolism through a starvation-induced autoregulatory loop. Nat Cell Biol, 2013. 15(6): p. 647–58.

13. Settembre, C., et al., TFEB links autophagy to lysosomal biogenesis. Science, 2011. 332(6036): p. 1429–33.

14. Martina, J.A., et al., MTORC1 functions as a transcriptional regulator of autophagy by preventing nuclear transport of TFEB. Autophagy, 2012. 8(6): p. 903–14.

15. Martina, J.A., et al., The nutrient-responsive transcription factor TFE3 promotes autophagy, lysosomal biogenesis, and clearance of cellular debris. Sci Signal, 2014. 7(309): p. ra9.

16. Roczniak-Ferguson, A., et al., The transcription factor TFEB links mTORC1 signaling to transcriptional control of lysosome homeostasis. Sci Signal, 2012. 5(228): p. ra42.

17. Settembre, C., et al., A lysosome-to-nucleus signalling mechanism senses and regulates the lysosome via mTOR and TFEB. EMBO J, 2012. 31(5): p. 1095–108.

18. Pastore, N., et al., TFEB and TFE3 cooperate in the regulation of the innate immune response in activated macrophages. Autophagy, 2016. 12(8): p. 1240–58.

19. Samie, M. and P. Cresswell, The transcription factor TFEB acts as a molecular switch that regulates exogenous antigen-presentation pathways. Nat Immunol, 2015. 16(7): p. 729–36.

20. Chen, H.D., et al., HLH-30/TFEB-mediated autophagy functions in a cell-autonomous manner for epithelium intrinsic cellular defense against bacterial pore-forming toxin in C. elegans. Autophagy, 2017. 13(2): p. 371–385.

21. Najibi, M., et al., An Evolutionarily Conserved PLC-PKD-TFEB Pathway for Host Defense. Cell Rep, 2016. 15(8): p. 1728–42.

22. Baba, M., et al., Folliculin encoded by the BHD gene interacts with a binding protein, FNIP1, and AMPK, and is involved in AMPK and mTOR signaling. Proc Natl Acad Sci U S A, 2006. 103(42): p. 15552–7.

23. Takagi, Y., et al., Interaction of folliculin (Birt-Hogg-Dube gene product) with a novel Fnip1-like (FnipL/Fnip2) protein. Oncogene, 2008. 27(40): p. 5339–47.

24. Tee, A.R. and A. Pause, Birt-Hogg-Dube: tumour suppressor function and signalling dynamics central to folliculin. Fam Cancer, 2013. 12(3): p. 367–72.

25. Hardie, D.G., AMPK: positive and negative regulation, and its role in whole-body energy homeostasis. Curr Opin Cell Biol, 2015. 33: p. 1–7.

26. Hardie, D.G. and M.L. Ashford, AMPK: regulating energy balance at the cellular and whole body levels. Physiology (Bethesda), 2014. 29(2): p. 99–107.

27. Hardie, D.G., F.A. Ross, and S.A. Hawley, AMPK: a nutrient and energy sensor that maintains energy homeostasis. Nat Rev Mol Cell Biol, 2012. 13(4): p. 251–62.

28. Possik, E., et al., FLCN and AMPK Confer Resistance to Hyperosmotic Stress via Remodeling of Glycogen Stores. PLoS Genet, 2015. 11(10): p. e1005520.

29. Possik, E., et al., Folliculin regulates ampk-dependent autophagy and metabolic stress survival. PLoS Genet, 2014. 10(4): p. e1004273.

30. Possik, E. and A. Pause, Glycogen: A must have storage to survive stressful emergencies. Worm, 2016. 5(2): p. e1156831.

31. Yan, M., et al., The tumor suppressor folliculin regulates AMPK-dependent metabolic transformation. J Clin Invest, 2014. 124(6): p. 2640–50.

32. Blagih, J., et al., The energy sensor AMPK regulates T cell metabolic adaptation and effector responses in vivo. Immunity, 2015. 42(1): p. 41–54.

33. Prantner, D., D.J. Perkins, and S.N. Vogel, AMP-activated Kinase (AMPK) Promotes Innate Immunity and Antiviral Defense through Modulation of Stimulator of Interferon Genes (STING) Signaling. J Biol Chem, 2017. 292(1): p. 292–304.

34. Possik, E. and A. Pause, Measuring oxidative stress resistance of Caenorhabditis elegans in 96- well microtiter plates. J Vis Exp, 2015(99): p. e52746.

35. Rohlfing, A.K., et al., Genetic and physiological activation of osmosensitive gene expression mimics transcriptional signatures of pathogen infection in C. elegans. PLoS One, 2010. 5(2): p. e9010.

36. Irazoqui, J.E., et al., Distinct pathogenesis and host responses during infection of C. elegans by P. aeruginosa and S. aureus. PLoS Pathog, 2010. 6: p. e1000982.

37. Troemel, E.R., et al., p38 MAPK regulates expression of immune response genes and contributes to longevity in C. elegans. PLoS Genet, 2006. 2(11): p. e183.

38. Chen, L., et al., AMPK activation by GSK621 inhibits human melanoma cells in vitro and in vivo. Biochem Biophys Res Commun, 2016. 480(4): p. 515–521.

39. Jiang, H., et al., GSK621 Targets Glioma Cells via Activating AMP-Activated Protein Kinase Signalings. PLoS One, 2016. 11(8): p. e0161017.

40. Yan, M., et al., Chronic AMPK activation via loss of FLCN induces functional beige adipose tissue through PGC-1alpha/ERRalpha. Genes Dev, 2016. 30(9): p. 1034–46.

41. Hong, S.B., et al., Inactivation of the FLCN tumor suppressor gene induces TFE3 transcriptional activity by increasing its nuclear localization. PLoS One, 2010. 5(12): p. e15793.

42. Betschinger, J., et al., Exit from pluripotency is gated by intracellular redistribution of the bHLH transcription factor Tfe3. Cell, 2013. 153(2): p. 335–47.

43. Martina, J.A. and R. Puertollano, Rag GTPases mediate amino acid-dependent recruitment of TFEB and MITF to lysosomes. J Cell Biol, 2013. 200(4): p. 475–91.

44. Petit, C.S., A. Roczniak-Ferguson, and S.M. Ferguson, Recruitment of folliculin to lysosomes supports the amino acid-dependent activation of Rag GTPases. J Cell Biol, 2013. 202(7): p. 1107–22.

45. Wada, S., et al., The tumor suppressor FLCN mediates an alternate mTOR pathway to regulate browning of adipose tissue. Genes Dev, 2016. 30(22): p. 2551–2564.

46. Efeyan, A., et al., Regulation of mTORC1 by the Rag GTPases is necessary for neonatal autophagy and survival. Nature, 2013. 493(7434): p. 679–83.

47. Tsun, Z.Y., et al., The folliculin tumor suppressor is a GAP for the RagC/D GTPases that signal amino acid levels to mTORC1. Mol Cell, 2013. 52(4): p. 495–505.

48. Peli-Gulli, M.P., et al., Feedback Inhibition of the Rag GTPase GAP Complex Lst4-Lst7 Safeguards TORC1 from Hyperactivation by Amino Acid Signals. Cell Rep, 2017. 20(2): p. 281–288.

49. Hasumi, Y., et al., Folliculin (Flcn) inactivation leads to murine cardiac hypertrophy through mTORC1 deregulation. Hum Mol Genet, 2014. 23(21): p. 5706–19.

50. Young, N.P., et al., AMPK governs lineage specification through Tfeb-dependent regulation of lysosomes. Genes Dev, 2016. 30(5): p. 535–52.

51. Engelmann, I., et al., A comprehensive analysis of gene expression changes provoked by bacterial and fungal infection in C. elegans. PLoS One, 2011. 6(5): p. e19055.

52. O’Rourke, D., et al., Genomic clusters, putative pathogen recognition molecules, and antimicrobial genes are induced by infection of C. elegans with M. nematophilum. Genome Res, 2006. 16(8): p. 1005–16.

53. Sinha, A., et al., System wide analysis of the evolution of innate immunity in the nematode model species Caenorhabditis elegans and Pristionchus pacificus. PLoS One, 2012. 7(9): p. e44255.

54. Ishii, K.J., et al., Host innate immune receptors and beyond: making sense of microbial infections. Cell Host Microbe, 2008. 3(6): p. 352–63.

55. Frisard, M.I., et al., Low levels of lipopolysaccharide modulate mitochondrial oxygen consumption in skeletal muscle. Metabolism, 2015. 64(3): p. 416–27.

56. Hansen, M.E., et al., Lipopolysaccharide Disrupts Mitochondrial Physiology in Skeletal Muscle via Disparate Effects on Sphingolipid Metabolism. Shock, 2015. 44(6): p. 585–92.

57. Kato, M., Site of action of lipid A on mitochondria. J Bacteriol, 1972. 112(1): p. 268–75.

58. McGivney, A. and S.G. Bradley, Action of bacterial lipopolysaccharide on the respiration of mouse liver mitochondria. Infect Immun, 1980. 27(1): p. 102–6.

59. Gordon, S., Phagocytosis: An Immunobiologic Process. Immunity, 2016. 44(3): p. 463–475.

60. Gray, M.A., et al., Phagocytosis Enhances Lysosomal and Bactericidal Properties by Activating the Transcription Factor TFEB. Curr Biol, 2016. 26(15): p. 1955–1964.

61. Schmidt, L.S. and W.M. Linehan, Molecular genetics and clinical features of Birt-Hogg-Dube syndrome. Nat Rev Urol, 2015. 12(10): p. 558–69.

62. Kauffman, E.C., et al., Molecular genetics and cellular features of TFE3 and TFEB fusion kidney cancers. Nat Rev Urol, 2014. 11(8): p. 465–75.

63. Karin, M., NF-kappaB as a critical link between inflammation and cancer. Cold Spring Harb Perspect Biol, 2009. 1(5): p. a000141.

64. Zhang, B.B., G. Zhou, and C. Li, AMPK: an emerging drug target for diabetes and the metabolic syndrome. Cell Metab, 2009. 9(5): p. 407–16.

65. Brenner, S., The genetics of Caenorhabditis elegans. Genetics, 1974. 77(1): p. 71–94.

66. Kamath, R.S., et al., Effectiveness of specific RNA-mediated interference through ingested double-stranded RNA in Caenorhabditis elegans. Genome Biology, 2001. 2(1).

67. Powell, J.R. and F.M. Ausubel, Models of Caenorhabditis elegans infection by bacterial and fungal pathogens. Methods Mol Biol, 2008. 415: p. 403–27.

68. Hoogewijs, D., et al., Selection and validation of a set of reliable reference genes for quantitative sod gene expression analysis in C. elegans. BMC Mol Biol, 2008. 9: p. 9.

69. Vincent, E.E., et al., Differential effects of AMPK agonists on cell growth and metabolism. Oncogene, 2015. 34(28): p. 3627–39.

